# Allopolyploidy expanded gene content but not pangenomic variation in the hexaploid oilseed *Camelina sativa*

**DOI:** 10.1101/2024.08.13.607619

**Authors:** Kevin A. Bird, Jordan R. Brock, Paul P. Grabowski, Avril M. Harder, Shengqiang Shu, Kerrie Barry, LoriBeth Boston, Christopher Daum, Jie Guo, Anna Lipzen, Rachel Walstead, Jane Grimwood, Jeremy Schmutz, Chaofu Lu, Luca Comai, John K. McKay, J. Chris Pires, Patrick P. Edger, John T. Lovell, Daniel J. Kliebenstein

## Abstract

Ancient whole-genome duplications (WGDs) are believed to facilitate novelty and adaptation by providing the raw fuel for new genes. However, it is unclear how recent WGDs may contribute to evolvability within recent polyploids. Hybridization accompanying some WGDs may combine divergent gene content among diploid species. Some theory and evidence suggest that polyploids have a greater accumulation and tolerance of gene presence-absence and genomic structural variation, but it is unclear to what extent either is true. To test how recent polyploidy may influence pangenomic variation, we sequenced, assembled, and annotated twelve complete, chromosome-scale genomes of *Camelina sativa*, an allohexaploid biofuel crop with three distinct subgenomes. Using pangenomic comparative analyses, we characterized gene presence-absence and genomic structural variation both within and between the subgenomes. We found over 75% of ortholog gene clusters are core in *Camelina sativa* and <10% of sequence space was affected by genomic structural rearrangements. In contrast, 19% of gene clusters were unique to one subgenome, and the majority of these were Camelina-specific (no ortholog in Arabidopsis). We identified an inversion that may contribute to vernalization requirements in winter-type Camelina, and an enrichment of Camelina-specific genes with enzymatic processes related to seed oil quality and Camelina’s unique glucosinolate profile. Genes related to these traits exhibited little presence-absence variation. Our results reveal minimal pangenomic variation in this species, and instead show how hybridization accompanied by WGD may benefit polyploids by merging diverged gene content of different species.

## Introduction

Whole-genome duplications (WGDs) are a near ubiquitous feature across the eukaryotic tree of life. When WGDs occur, they are believed to enable evolutionary novelty and adaptability through changes in functional genomic content via the retention and divergence of duplicate genes, which drive the expansion of gene families and the rewiring of genetic networks (Freeling and Thomas, 2006; Soltis and Soltis, 2016; De Smet et al. 2017). This is primarily exemplified by the presence of WGDs close to the origin of flowering plants and vertebrates (Wagner, 2008). In contrast to these clear long-term benefits, more recent WGDs have more ambiguous benefits as illustrated by their higher extinction and lower diversification compared to closely related diploids. This has led to an apparent paradox that in the short-term, polyploidy may often be an evolutionary dead end (Mayrose et al. 2011). Still, there is evidence that the masking effect of duplicate genes, WGD-induced transcriptomic and epigenomic variation, and short-term network effects of WGD may provide enhanced evolvability in newly formed polyploids, especially under environmental stress (Osborn et al. 2003; Leitch and Leitch, 2008; Van de Peer et al. 2021). A further possibility is that WGD promotes evolvability in newly formed polyploids by increasing genomic variation.

Gene duplication has long been recognized as a major source of genetic variation due to the redundancy of duplicated genes (Ohno, 1970; Wagner, 2008). This variation is not limited to single nucleotide polymorphisms; the subsequent duplication and loss of genes within a species create a vast amount of variation in functional gene content that contributes to the pangenome (the entire set of genomic sequences within a species). WGDs are an immediate source of a large pool of duplicate genes and so may promote evolvability in polyploids by generating greater pangenomic variation within a species. Recent work supports this contention. An analysis of a ploidy series in the genus *Cochlearia* identified an excess of large genomic structural variants (SVs) compared to diploids and a progressive effect as ploidy increased (Hamala et al. 2023). In *Brassica napus,* high levels of gene presence-absence variation (PAV) were observed with a substantial amount driven by homoeologous exchanges (non-reciprocal translocations between orthologous regions of chromosomes), which represent a distinct mechanism of pangenomic variation from that found in closely related diploid species (Hurgobin et al. 2018; Song et al. 2020; Bayer et al. 2021). The observations of unique mechanisms to generate pangenomic variation and a superior capacity to accumulate pangenomic variation in polyploids compared to diploids lead to the prediction that polyploid species will have large and dynamic pangenomes.

While polyploidy from both within-species WGD (autopolyploidy) or from the merger of two or more diverged lineages (allopolyploidy) can increase gene redundancy, allopolyploidy may have an additional mechanism to create novelty or variation. If the diverged lineages that hybridize to form an allopolyploid have diversified to possess unique gene content, the combination of the two genomes will expand the organisms’ genetic repertoire. Under this model, the allopolyploid progenitors diverged in gene content due to ongoing drift and selection that occurs specifically in each lineage. For example, when two species diverge from a common ancestor, they will begin with the approximately same gene content but over time develop their own unique gene content through these processes. If allopolyploidy merges these genomes, the new species combines all the genes from both progenitors and increases its functional gene space in comparison to the ancestral diploid species. For this model to be plausible there must be sufficient variation in gene content at the genus level. Studies on gene content variation of diploids at the genus level have shown that species have diverged in gene content to a significant extent (Qiao et al. 2021; Bozan et al. 2023; Li et al. 2023; Cochetel et al. 2023; Ferguson et al. 2024). Variable gene content distinguished the major clades of the petota group in potato, and a large number of clade-specific genes were observed (Bozan et al. 2023). Similarly, *Vitis* species showed high levels of species-specific genomic content (Cochetel et al. 2023). Further supporting this model, the genomes of *Brassica napus* (Chalhoub et al. 2014) and hexaploid wheat (IWGSC, 2014) reported an appreciable level of subgenome-specific genes. These studies establish that diploid species are frequently diverged in gene content, but the contribution of this divergence to polyploid genomes and its impact on polyploid evolution is still largely unknown.

To study WGD and how it may influence the genome evolution in a recent allopolyploid, we are using *Camelina sativa* (2n=6x=40). Camelina is a member of the *Brassicaceae* family and a notable oilseed crop with a hexaploid genome structure generated by the merger of three diploid genomes. There is renewed interest in the species for use in the bioeconomy as a source of renewable biofuel, plant factory for high-value molecules, and emerging model system for plant science research. This interest is from a blend of metabolic novelty in Camelina and key agronomic properties like relatively disease resistance (Séguin-Swartz et al., 2009), ability to grow in saline and/or marginal soils (Moser, 2010), and short generation time (Gugel and Falk, 2006). The initial Camelina reference genome confirmed the allopolyploid genome structure of Camelina with three distinct subgenomes (Kagale et al. 2014). Subsequent work identified the most likely progenitor species as the diploid *C. neglecta* (2n=2x=12), a putatively extinct *C. neglect-*like lineage with seven chromosomes (2n=2x=14), and *C. hispida* (2n=2x=14) (Fig 1A; Mandáková et al. 2019; Chaudhary et al. 2020; Mandáková and Lysak, 2022; Brock et al. 2022b). The hexaploid formation has been estimated at approximately 65,000 years ago (Brock et al. 2022b) and domestication occurred around 8,000 years ago in western Asia, likely in the Caucasus region (Brock et al. 2022a). The recent polyploid formation and domestication, as well as the detailed knowledge of progenitor species, make Camelina an ideal system to investigate gene content variation arising from recent WGDs.

Further boosting Camelina’s potential as a system to study the evolutionary consequences of WGD within allopolyploids is its phylogenetic proximity to the model plant Arabidopsis. Camelina is estimated to have diverged from Arabidopsis around 8-17 million years ago (Kagale et al. 2014; Hohman et al. 2015), and both species are in the tribe *Camelineae,* making it more closely related to Arabidopsis than species in the *Brassica* genus. This proximity makes Camelina especially amenable to mapping the functional impact of genomic variation. By exploiting the detailed functional characterization of genes in Arabidopsis, we can more precisely infer the function of Camelina genes, interpret their variation, and test hypotheses about the influence of recent WGD on short-term functional and evolutionary changes.

Key systems where Arabidopsis mechanistic information and Camelina agronomic properties overlap are flowering time, seed oil production and glucosinolate production. These traits all exhibit Camelina specific novelty which can be interrogated using extensive mechanistic information from Arabidopsis mechanistic information. The winter and spring-types in Camelina, differentiated by their requirement for vernalization prior to flowering, facilitate their use as cover crops or rapid-cycling rotational oil crops and allow comparison to the well-characterized vernalization pathway in Arabidopsis (Choi et al. 2015). The oil-rich seeds of Camelina are of greatest interest for the bioeconomy (Gugel and Falk, 2006; Berti et al. 2016; Zanetti et al. 2017), and the detailed characterization of lipid metabolism in Arabidopsis (Li-Besson et al. 2010) provides the foundation to explore the evolution of Camelina’s oil profile, notable for high levels of unsaturated and omega-3 fatty acids that may meet different end user needs. Like other members of the *Brassicales*, Camelina produces glucosinolates, a class of sulfur-rich, amino acid-derived specialized metabolites with a role in environmental adaptation to a multitude of biotic and abiotic stresses. Most *Brassicaceae,* like Arabidopsis and Brassica, have shorter chain aliphatic glucosinolates where the methionine precursor has two to seven carbons added leading to a final aliphalitc glucosinolate of between three (3C) and eight carbons (8C). In contrast *Camelina sativa* and closely related *Brassicaceae* in the *Capsella* and *Neslia* genera have increased this chain length to create glucosinolates with nine (9C), ten (10C) or eleven (11C) glucosinolates (Fahey, 2001; Czerniawski et al. 2021). The characterization of the causal basis of glucosinolate variation in Arabidopsis (Sonderby et al. 2010) can aid in identifying changes responsible for Camelina’s unique glucosinolate profile. These pathways provide an opportunity to query genome sequences for a better understanding of how genomic variation from WGD relates to metabolic and phenological changes.

## Results

### Genome assembly and annotation

**Fig 1.**
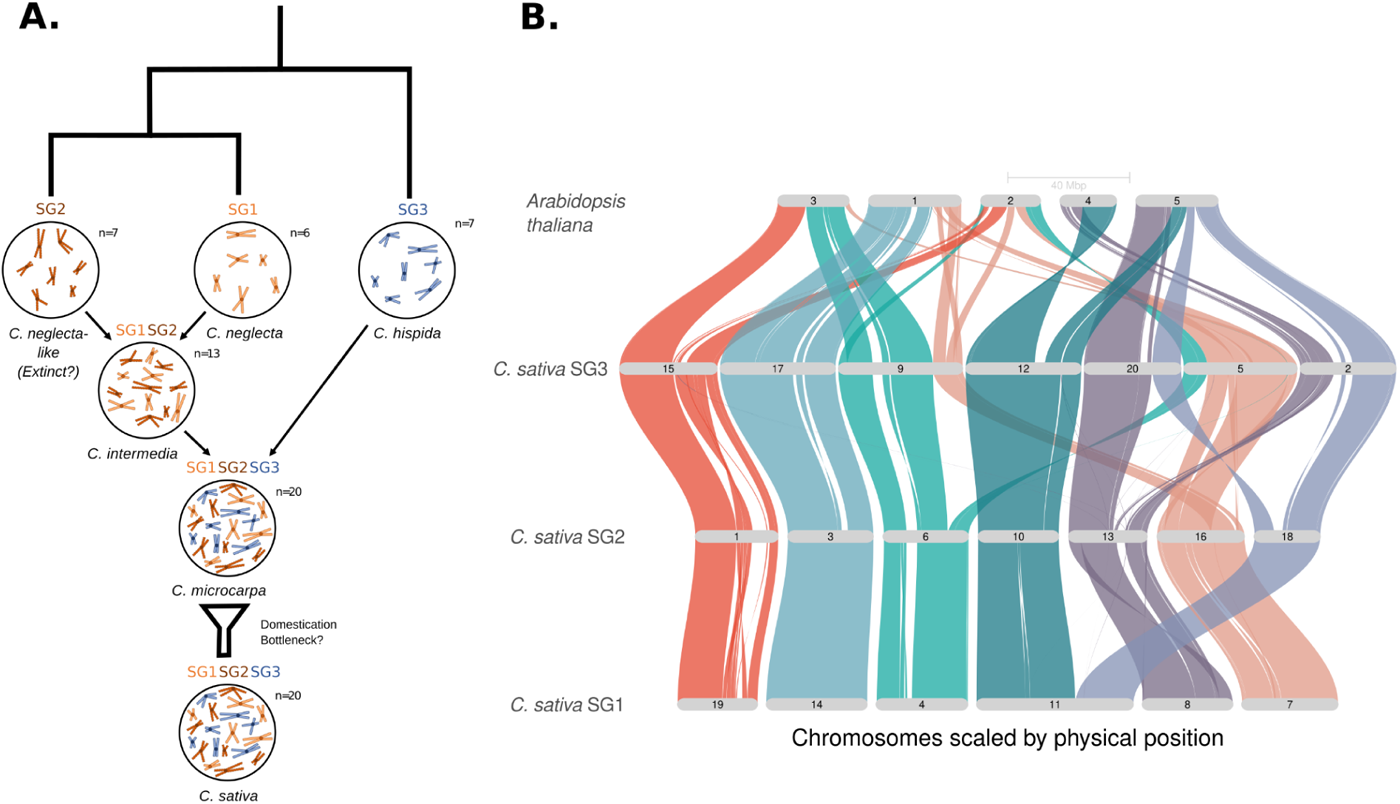
**A)** Summary of *Camelina sativa* progenitors and allopolyploid formation. **B)** Synteny between *Arabidopsis thaliana* and the three *C. sativa* subgenomes from the reference assembly Prytzh.

Here we present twelve complete, chromosome-scale de novo assemblies of diverse *Camelina sativa* accessions. Comparative pangenomic analyses identified core and dispensable gene clusters, generated a catalog of structural variants, and allowed gene orthgroup partitioning between species and individual subgenomes. This showed substantial gene content unique to subgenomes and to Camelina, suggesting allopolyploidy increased the functional gene content of the hexaploid Camelina in comparison to the diploids. In contrast, the gene content and genome structure were highly conserved within the species, resulting in low levels of pangenomic variation. Dissecting specific metabolic pathways showed that the majority of novelty within Camelina appears to be created by orthogroup variation between Arabidopsis and Camelina at the pathway level.

**Fig 2.**
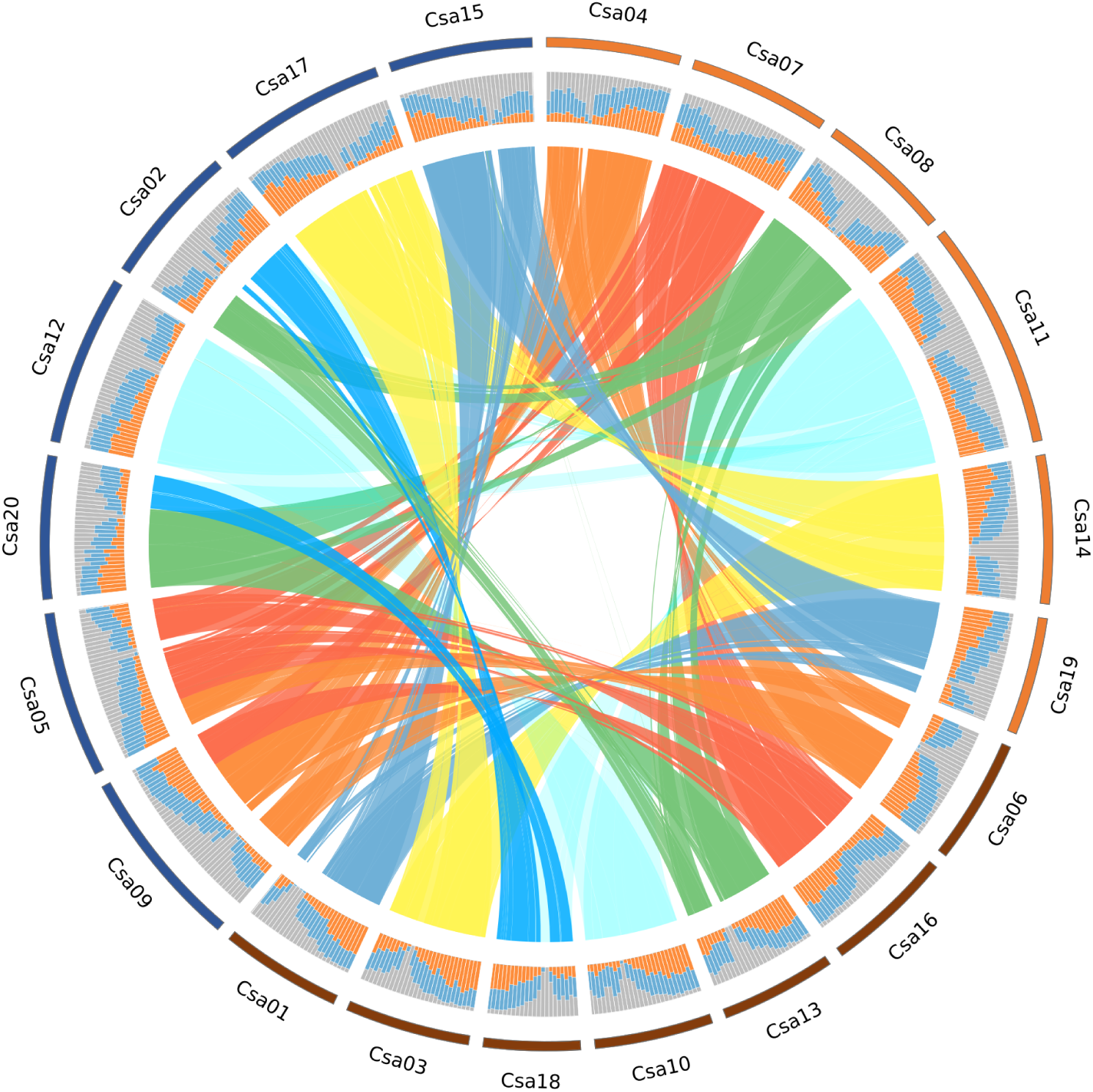
Circos plot of reference assembly Prytzh, outer ring colors represent the distinct subgenomes, middle ring shows % of genic (orange), repetitive (blue) and unannotated (grey) sequence in 1Mb windows, and ribbons in the middle indicate syntenic regions between subgenomes based on GENESPACE results.

We selected twelve Camelina lines for sequencing, representing the diversity within the current USDA collection. These include three winter types (Joelle, Acsn 226, CN113691) and nine spring types including lines derived from breeding programs (ASCN226, Borowska, Prytzh, Svalof, CN113691), elite or improved cultivars (Lindo, Licalla), and “wild” accessions (CAM70, CAM116) collected from Western, Central, and Eastern Europe, the geographic regions representing the major population structure divisions in the collection (Li et al. 2021).

To generate the genome assemblies, we employed a whole-genome shotgun strategy using PacBio circular consensus sequencing (CCS) HiFi and Illumina platforms. For Prytzh, we coassembled PacBio HiFi (83.51X) and Hi-C reads using HiFiAsm+HIC. We then scaffolded the contigs using the JUICER pipeline to generate a 639.1Mb main genome assembly, which was then short-read polished using 2×150 Illumina reads (65.1X) to correct homozygous SNPs and short indels. The final Prytzh assembly contains 20 scaffolds representing the 20 chromosomes from the three subgenomes (Fig 1B, Fig 2). Chromosomes from SG1 and SG2 are largely colinear, with Chr10 and Chr18 from SG2 (7 chromosomes) aligning with opposite sides of Chr11 from SG1 (6 chromosomes; Fig 1B; Fig 2), while there have been several chromosomal rearrangements between SG3 and SG1/SG2 (Fig 1B). These recapitulate observations from cytogenetic and comparative genomic work in *C. sativa* and the diploid progenitor genomes (Mandakova et al. 2019; Martin et al. 2022).

From the Pryzth assembly, we generated 20,563 unique, non-repetitive, non-overlapping 1Kb syntenic markers to assist the assembly of the other genotypes. For the other eleven genotypes, we assembled PacBio HiFi (19.54-69.43X) reads using HiFiAsm and used the syntenic markers from Pryzth to correct contig breaks and to orient, order, and join the contigs into chromosome-scale scaffolds. The scaffolds were then short-read polished using 2×150 Illumina reads (48.4-63.0X). In total, we generated twelve assemblies ranging in size from 598-656Mb, with 98.2%-100% of sequence present in chromosome-scale scaffolds representing the 20 *C.sativa* chromosomes (Table 1). Each assembly is highly contiguous, constructed from 26-95 contigs, with nine assemblies containing at least one chromosome assembled from a single contig. Additionally, we sequenced 26-32 telomeres per assembly. Based on BUSCO evaluation, the completeness of the twelve genomes each exceeded 99%. Overall, all twelve assemblies represent a great improvement in comparison to the existing reference genome, DH55 (Kagale et al. 2014).

We generated well-supported gene model annotations to accompany each of the twelve genome assemblies, using a two-pass method to leverage the information from the pangenome to improve the gene model predictions for each genotype. The number of protein-coding genes ranged from 69,153 to 74,852 (Table 1), which is notably lower than the 89,418 genes in the annotation for the DH55 reference genome (Kagale et al. 2014). Gene density in the Camelina genome tended to be higher in the chromosome arms and was often asymmetric, showing much greater density on one arm compared to the other. The patterns of gene density were broadly similar across syntenic chromosomes (Fig 2, Fig S1).

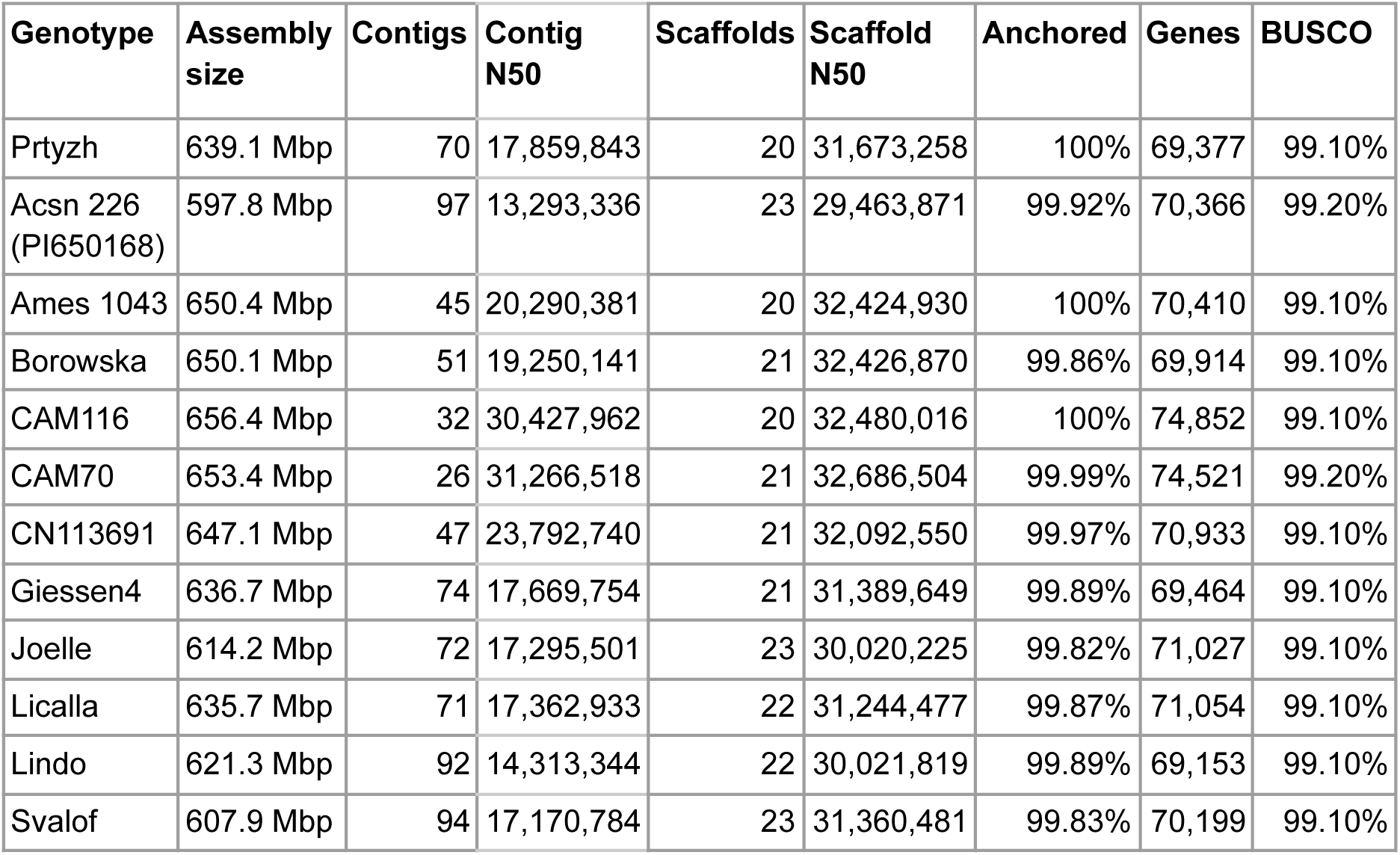

### Pangenome graph reveals a small dispensable genome

We generated a pangenome graph with Minigraph-Cactus using Prytzh as the backbone. The final graph is 742Mb and made from 11.3M nodes and 15.3M edges (Table S1). A high amount of sequence (93.1%-98.6%) from each assembly is retained in the final graph (Table S2). The core graph, containing sequence shared by all haplotypes, is 526Mb (70.9%), and 63.7Mb (8.6%) of the graph is unique sequence found in only single haplotypes (Table S1). The unaligned sequence from each assembly is clipped out of the final graph, and this unaligned sequence is comprised of 61.3%-92.4% repeat content and very little genic sequence (Table S2). There is the beginning of a plateau in the amount of new sequence incorporated with additional haplotypes, indicating the graph is approaching a closed graph and contains much of the alignable, non-repetitive sequence in Camelina (Fig 3).

**Fig 3.**
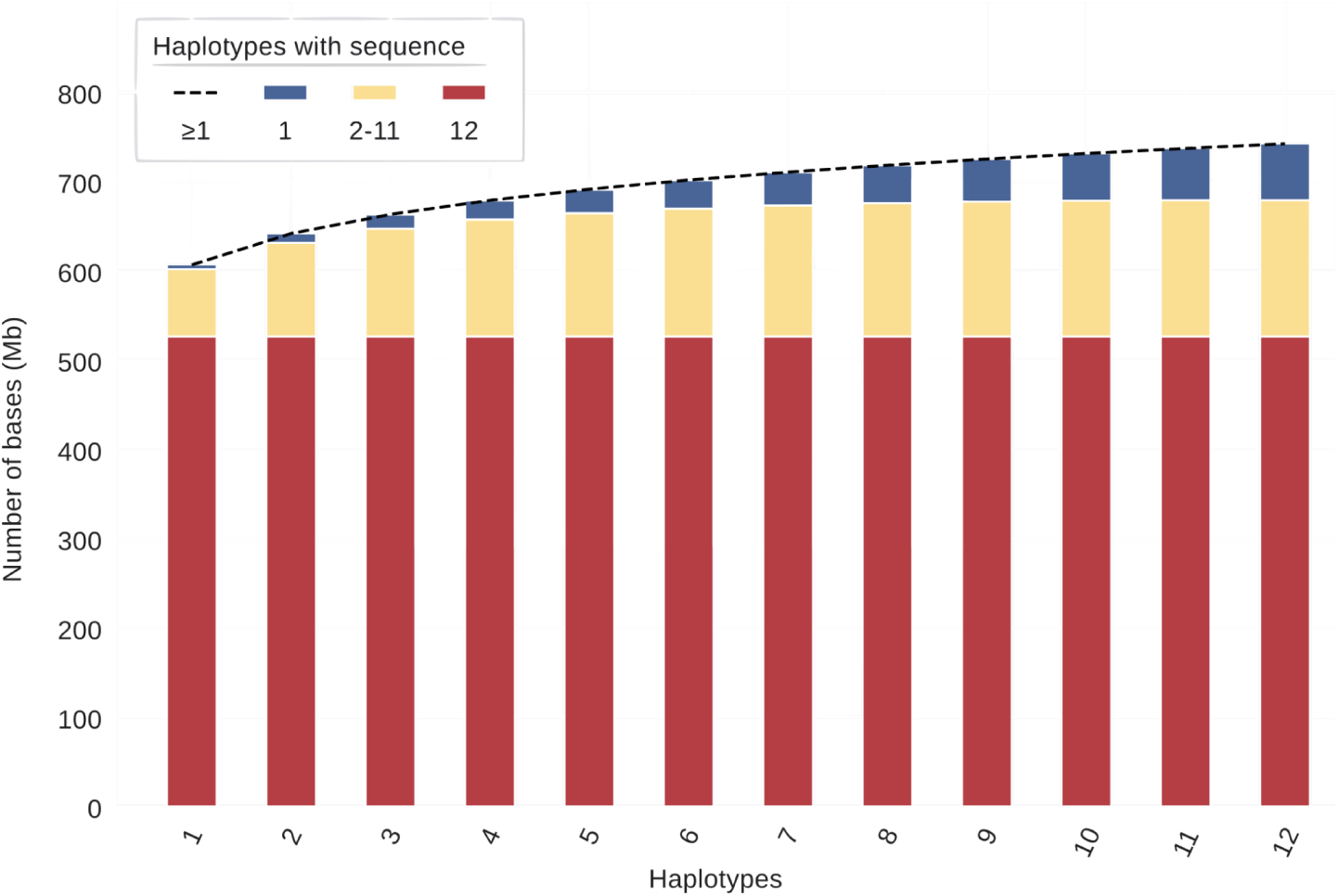
Amount of sequence in the genome graph that is shared by different numbers of haplotypes (assemblies). Figure only represents sequence in the final graph and does not contain sequences “clipped out”.

### Camelina shows low levels of gene presence-absence variation

To understand how the pangenome is structured in *Camelina sativa*, we determined pangenomic variation at the gene-family level using phylogenetic orthology inference by applying Orthofinder2 with *Arabidopsis thaliana* as the outgroup. This identified 30,327 hierarchical orthogroups that contained Camelina genes (Table S3-S4. Of these, 22,967 (76%) were present in all twelve genomes and classified as core orthogroups; an additional 5,683 (19%) were present in 2-11 genomes and classified as dispensable; and 1,567 (5%) were present in only a single genome and classified as unique (Fig 4A; Table S5). This level of gene PAV is less than observed in *B. napus* where 56% of genes clusters were core (Song et al. 2020) and similar to hexaploid wheat where 74% of genes were core (Walkowiak et al. 2020). The size of the Camelina pangenome increased as more genomes were added and did not reach an asymptote. Thus, incorporating additional genomes will identify additional unique and dispensable genes (Fig 4B).

**Fig 4.**
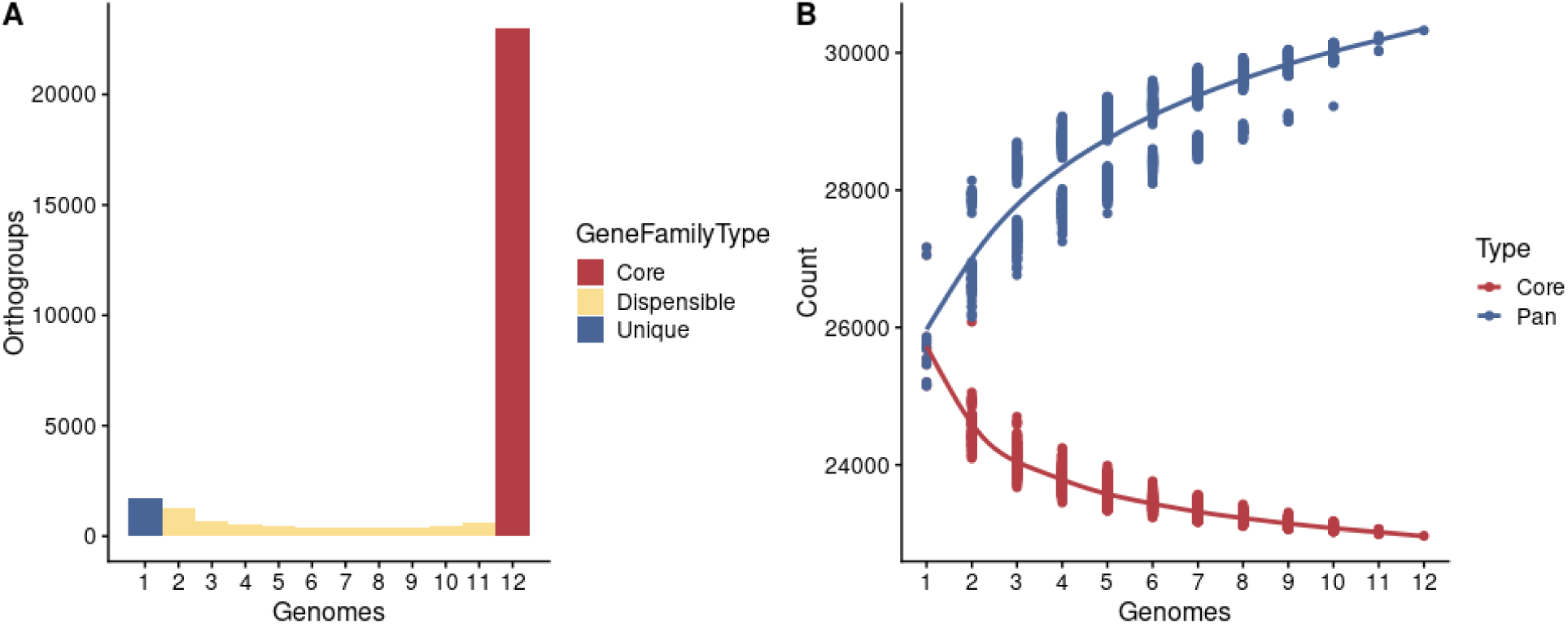
Orthogroup presence-absence variation. **A)** Breakdown of core, dispensable, and unique orthogroup content among our twelve genomes. **B)** Core and pan genome modeling for Camelina accessions. Each point represents the size of core and pangenome content for every possible combination of that sample size from our genomes and the line represents a fit loess model. The bimodal distribution in the pangenome content at less than twelve genomes is caused by genotypes CAM70 and CAM116 being highly similar and their presence or absence creating this bifurcation. Note the y-axis does not start at 0.

### Transposable elements contribute to genomic differences and show distinct patterns of presence-absence variation

To investigate how polyploidy may influence TE variation, we queried the TE content in the twelve Camelina genomes. This showed that the vast majority of TE families are core with the remaining dispensable TEs present in low numbers and without structure between pairs or larger subgroups of individuals (Fig S2-S3). Among annotated TEs, slightly more TE families were found to be core (49.7%) compared to dispensable (48.8%), and a small fraction were unique to individual genomes (1.5%) (Fig 5A).

**Fig 5.**
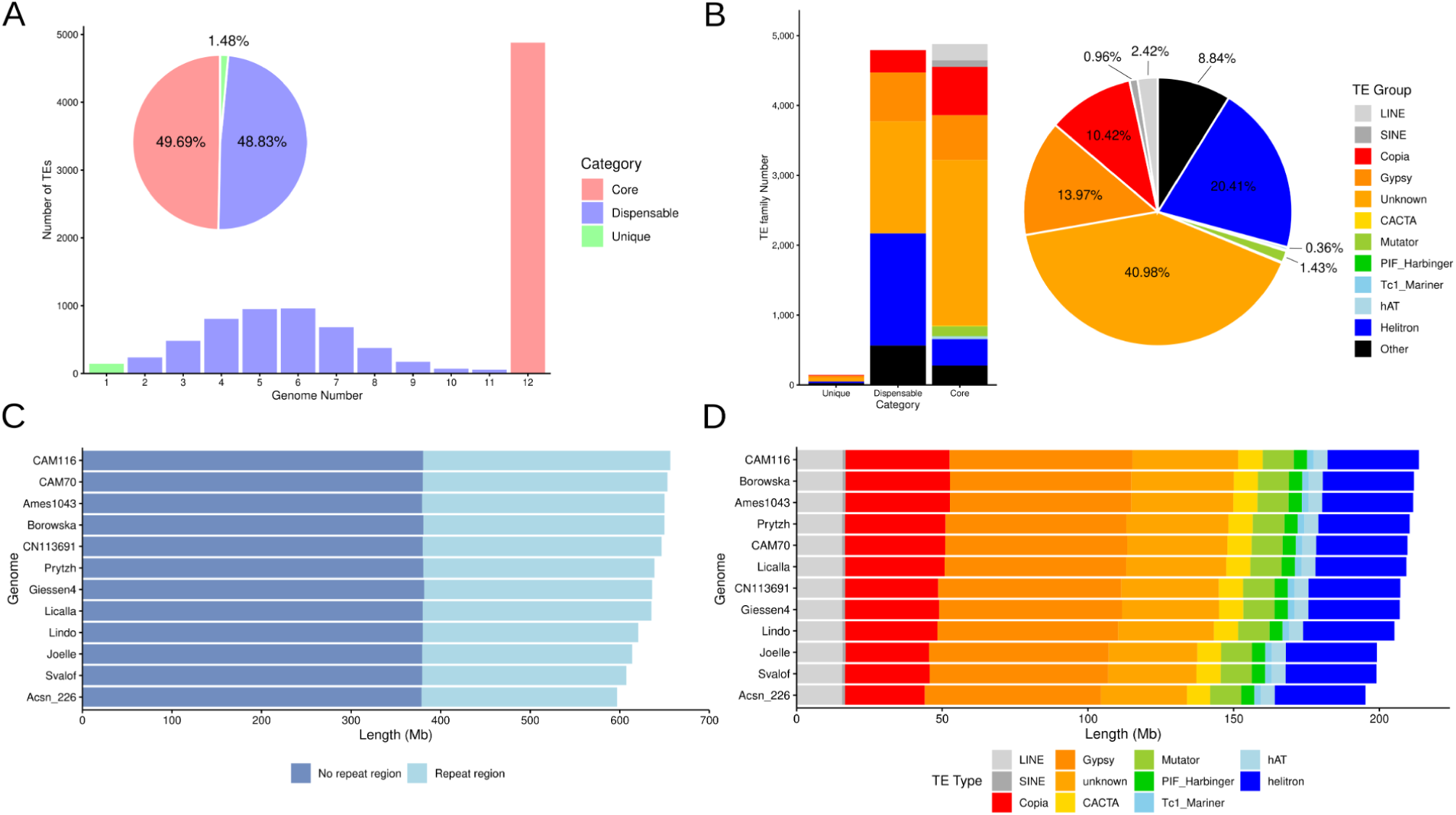
Results of the pangenome annotation of TEs across twelve *C. sativa* genomes. **A)** Number of TE families which were found to be unique to a single individual, dispensable (found in between 2 – 11 genomes), and core (found in all twelve genomes). **B)** Breakdown of the number of each TE type across the entire pangenome for unique, dispensable, and core categories. **C)** Proportion of each genome which was found to contain no repeat regions and repeat regions. **D)** Proportion of each TE class within the repeat fraction of the genome, colored by TE type.

The set of variable TE families (those TEs present in 2 – 11 genomes) showed a nearly normal distribution in content which contrasts with the heavy skew towards core TE families seen in other species like Arabidopsis (Fig 5A) (Kang et al. 2023). This also differs from the presence-absence variation in orthogroups that was heavily skewed to core. We find little structure in TE families among the individuals in the pangenome, such that the lack of population structure is likely a combination of the minimal level of genetic differentiation between Camelina accessions and potential inter-crossing amongst genotypes. Among those TEs annotated, LINEs, SINEs, and unknown TEs were almost exclusively core, whereas DNA and LTR elements represented the majority of unique and dispensable TEs (Fig 5B). The size of the repeat proportion of genomes ranged from 250 Mb (CAM116) to 193 Mb (Acsn 226) which largely aligns with differences in assembly size, whereas the proportion of the non-repeat regions of the genomes assembled were highly consistent (Figure 5C). This variation in repeat content was primarily driven by LTR Copia elements (Figure 5D).

### *Camelina sativa* has a highly stable genome structure

After assessing the gene and TE level variation we proceeded to investigate structural variation amongst the twelve genomes. Analysis of macrosynteny using GENESPACE (v.1.2.3; Lovell et al. 2021) revealed extensive conserved genome structure across the twelve genomes with most chromosomes exhibiting no large structural rearrangements (Fig 6; Fig S4-S5). We further identified structural variants in the pangenome with pairwise whole-genome alignments against a common reference, Prytzh, using the program SyRi (v1.6.3; Goel et al. 2019). This showed that the genome structure is highly consistent, with 573.9-616.7Mb (on average 94% of the genome) being syntenic and no evidence for large structural variation. In contrast, 16.3-65.4Mb (On average 6%) showed evidence of structural rearrangements. (Fig 7A; Table S6). In terms of structural variation, we identified 1.4-10.0Mb of inverted regions and 0.025-0.81Mb of translocated regions. We queried for inter-subgenomic translocations that may represent segregating homoeologous exchange events, and only detected 2-5 per line, representing no more than 300kb (Table S7). In contrast, the vast majority of SVs were simple loss or gain of sequences in relation to the reference. Approximately 25.9-47.3Mb were lost per genome in comparison to the reference by a combination of deletions, copy loss, and regions in the reference that did not align to the query genome. A slightly higher level of sequences was gained per-genome in comparison to the reference with 17.9-63.7Mb coming from a combination of insertions, copy gain, and sequences in the query genomes that did not align to the reference genome; Fig 7B; Table S8). The length of these SVs ranged from 2bp for indels to Mb scale variants including a 2.8Mb Inversion in CAM116, a 2.4Mb Inversion in Joelle, and several sequence gains or losses novel to query genomes on the scale of 4-5Mbs (Fig 7C; Table S9). When large SVs were broken down by subgenome, we observed that SG3 exhibited the largest number of SVs, driven primarily by a greater number of inversions, and copy number changes (Fig7D; Table S10). Previous analyses have identified that SG3 is slightly dominant in terms of expression (Kagale et al. 2014; Kagale et al. 2016).

**Fig 6.**
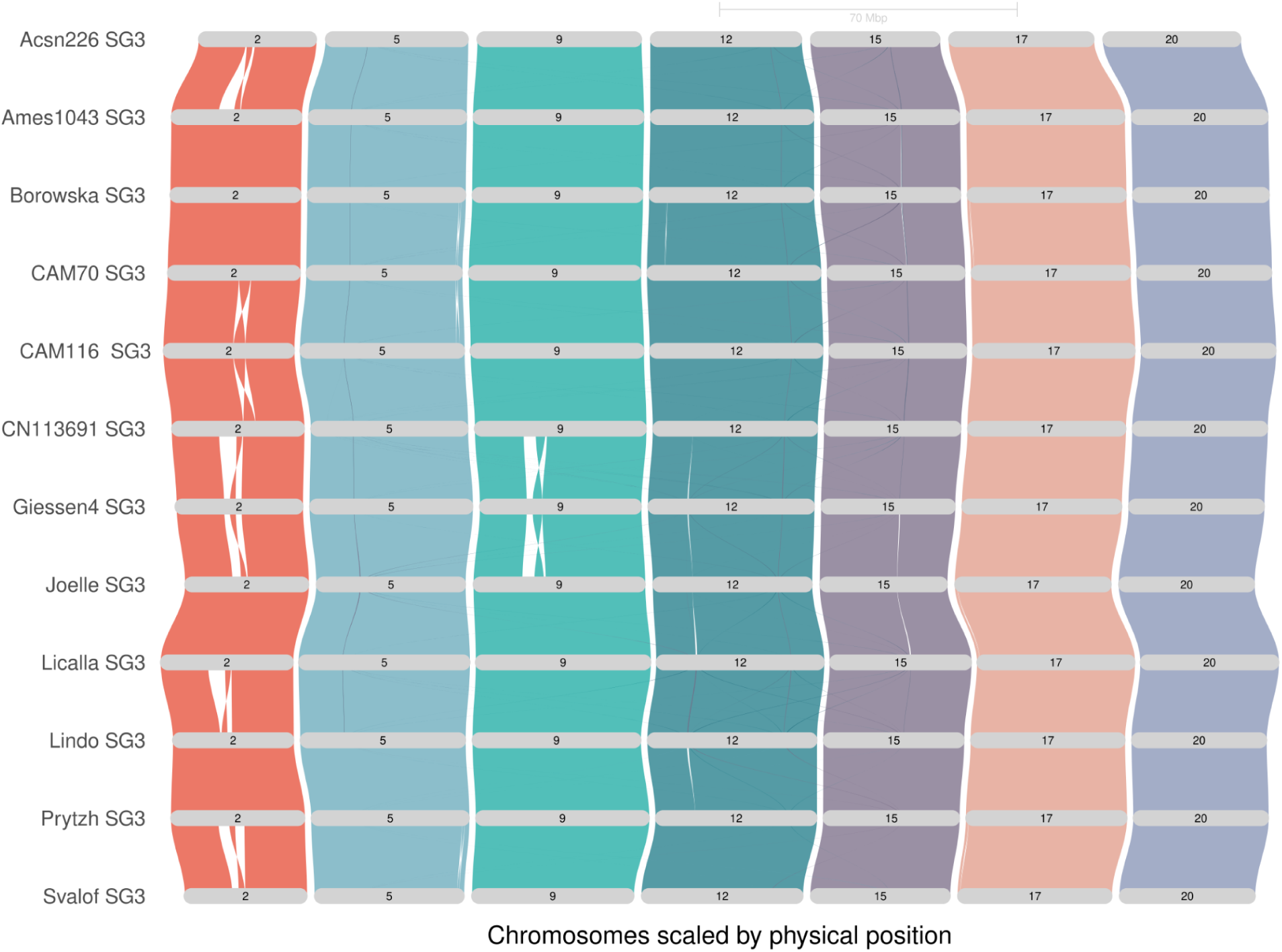
Riparian plot of macrosynteny for *C. sativa* subgenome 3 across all twelve assemblies.

**Fig 7.**
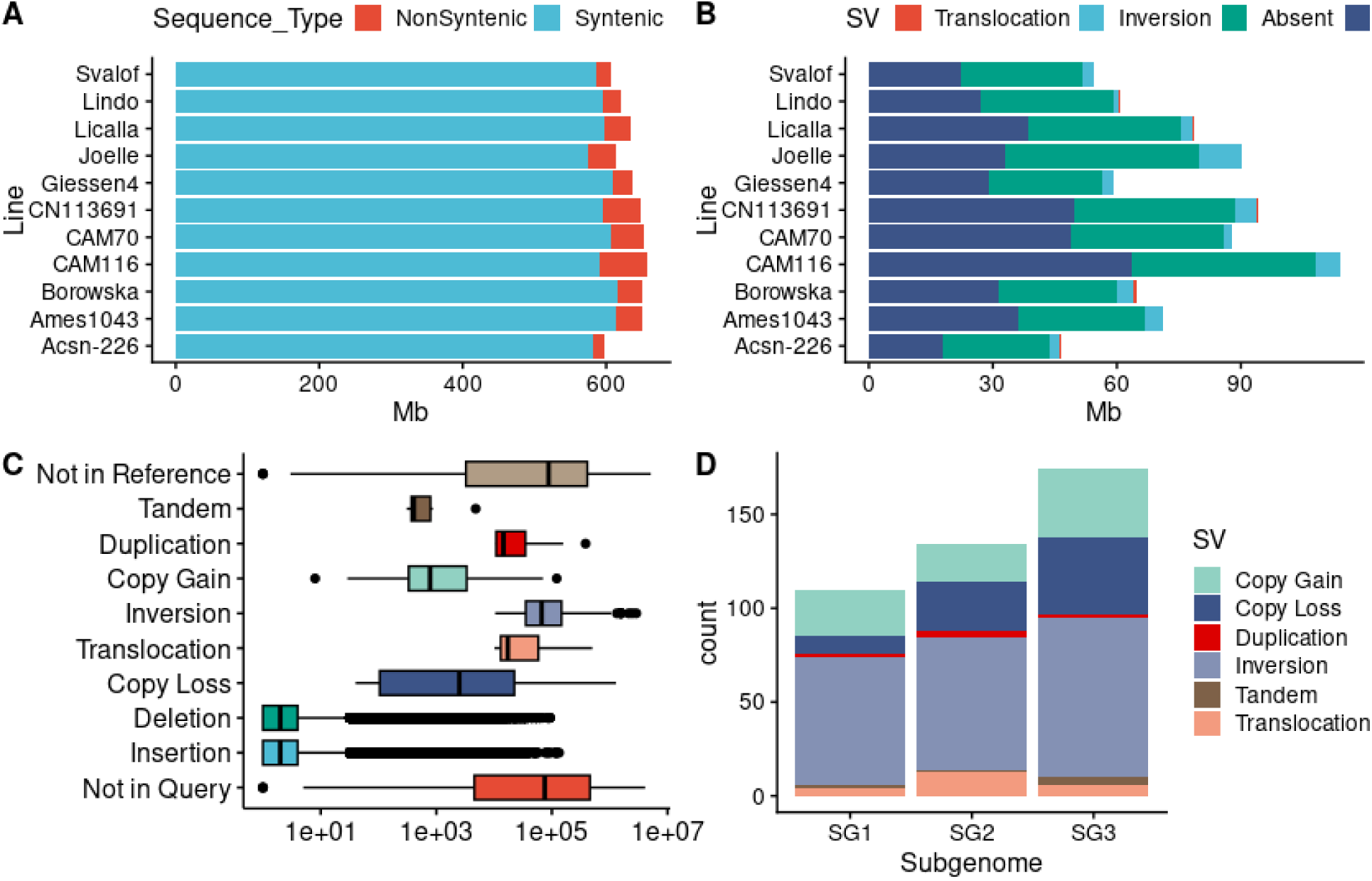
Results of the structural variant identification from comparison of eleven *C. sativa* genomes with the Prytzh assembly. **A)** Size in Mb of genomic regions syntenic between assemblies and non-syntenic due to structural variation. **B)** Size in Mb of non-syntenic regions of the genomes, broken down by SV type. **C)** Distribution of SV size, in bp, for all SV classes of interest. **D)** Number of large SVs by subgenome, does not include indels.

### Allopolyploidy integrates divergent gene content from progenitors

Whole-genome duplication is a key driver of evolutionary innovation with a focus on how the duplication of genes can facilitate their subsequent functional diversification. However, these processes can be countered by pangenome variation that can result in the loss of genes over time. Meanwhile, the ways in which pangenome variation influences gene content variation across species, and its impact on polyploid function, remain less studied. Camelina, by being an allopolyploid, allows us to query the potential for gene content divergence to occur over time by comparing the different subgenomes to each other and to the *Arabidopsis thaliana* genome. Further, it is possible that by merging three different genomes, an allopolyploidy event may further contribute to evolution by merging unique genomic material from the progenitors. We sought to explore the impact of pangenome partitioning on the Camelina genome by comparing the presence-absence variation of hierarchical orthogroups between Arabidopsis and the three Camelina subgenomes (Table S11).

We found that of the 32,109 hierarchical orthogroups identified by Orthofinder across the Camelina and Arabidopsis genomes, 7,698 (24%) did not have a corresponding ortholog in Arabidopsis. That suggests that these represent either genes lost in the reference Arabidopsis genome Col-0, but maintained in Camelina, or novel orthogroup gains in the Camelina lineage (Fig 8A; Table S12). An additional 1,782 (6%) were present in Arabidopsis, but did not have any orthologs across the Camelina genomes. Dissecting this further to subgenome-specific gene content within Camelina, we identified that 21,646 (64% of all orthogroups) were present at least once across all 3 subgenomes. There is also extensive partitioning of subgenome orthogroup content with 6,204 (19%) being subgenome-specific orthogroups. The most (2,523) unique to subgenome 3, the least (1649) unique to subgenome 2, and 2,042 unique to subgenome 1 (Fig 8A; Table S12). Thus, the allopolyploidy event leading to Camelina functionally increased the orthogroup content by merging genomes that had different orthogroups.

To better understand the functionalities associated with the subgenome-specific orthogroups we explored GO term enrichment among these genes. We found that orthogroups unique to subgenome 1 were enriched for several biological processes. The strongest enrichment was for Seryl-tRNA aminoacylation while also including RNA methylation, transport, and splicing, auxin response and signal transduction (Fig 8B; Table S13). Orthogroups unique to subgenome 2 were most strongly enriched for catabolism and surveillance of mRNA, and included terms for alanyl and aspartyl tRNA aminoacylation, carbon fixation, small GTPase mediated signal transduction, and phospholipid biosynthesis (Fig 8C; Table S14). Subgenome 3 unique orthogroups were most enriched for protein polymerization and mRNA catabolism, and included several other processes including defense response, response to auxin, and response to biotic stimulus. We also observed an enrichment for several microtubule-based GO terms, primary metabolic traits like sucrose, glutamate, and aromatic amino acid biosynthesis, and tRNA aminoacylation of glutaminyl and alanyl (Fig 8D; Table S15). This suggests that the subgenomes may have contributed new functionality to the polyploid upon merging.

**Fig 8.**
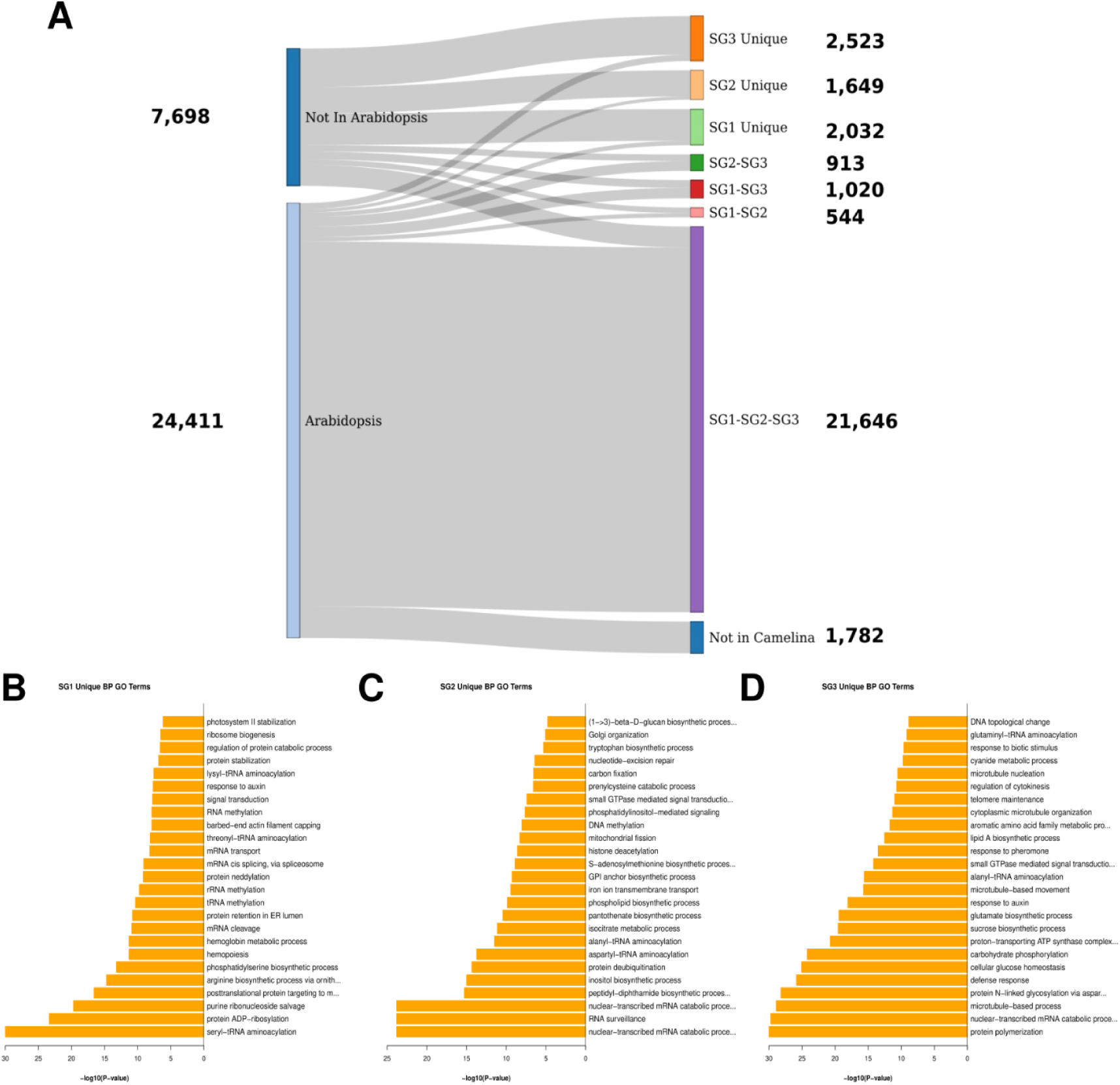
**A)** Breakdown of species and subgenome specific orthogroups. **B)** Top 25 Biological Process GO terms for genes unique to subgenome 1. **C)** Top 25 Biological Process GO terms for genes unique to subgenome 2. **D)**Top 25 Biological Process GO terms for genes unique to subgenome 3.

### An inversion shared by spring-type varieties may relate to vernalization requirement

A key agronomic trait in *Camelina sativa* is the variation exhibited between winter types that require vernalization before the transition from vegetative growth to floral initiation and spring types that do not. To investigate the potential utility of these genome sequences in mediating phenotypes in Camelina, we first focused on flowering time variation. Winter habits appear to be the ancestral state as the *Camelina* diploid species require vernalization. Our selected genomes include three individuals that are winter types and nine spring types. We combined these genome sequences with the information from the well-characterized molecular pathways controlling flowering and vernalization requirements in *Arabidopsis thaliana* to explore if any pangenome variation may influence the genetic basis of vernalization requirement in *C. sativa*. We found that all sampled Camelina lines contain orthologs of FLC, the Frigida complex (FRI, FRL1, FLX, FES1, SUF4), and Frigida-like 2, which have been shown in Arabidopsis to control natural variation in vernalization requirement (Michaels et al. 2004; Shindo et al. 2005; Choi et al. 2011). All of these genes were maintained in triplicate, except for Frigida-Like2 which contained two copies across the twlve genomes and FLC expressor (FLX), which contained two copies in eleven lines, and three copies in CN113691, a winter type. Thus, there was no link of gene level PAV to vernalization in this data.

Since gene PAV in known core regulators of vernalization requirement did not obviously correspond to winter and spring habit, we explored structural variation. Querying structural variants for association with vernalization showed that a large (∼100kb) polymorphic inversion on chromosome 11 had one orientation in all spring lines with a different orientation in all winter lines (Fig 9). The spring orientation of this structural variation is different from the same region in the other two subgenomes, suggesting the spring line orientation of this SV is the derived state. The inverted region contained approximately 90 genes, including a Frigida-like gene orthologous to FRL4-B (AT4G14900). This region also includes an ortholog of the circadian gene Time For Coffee (TIC), an ortholog of the Far Red Impaired Response 1 (FAR1) gene, and a tandem array of early-light induced protein (ELIP) genes. While this suggests that this SV may influence flowering time by altering one or more of these genes, further work will be necessary to see if this region and any of the contained genes influence vernalization requirement.

**Fig 9.**
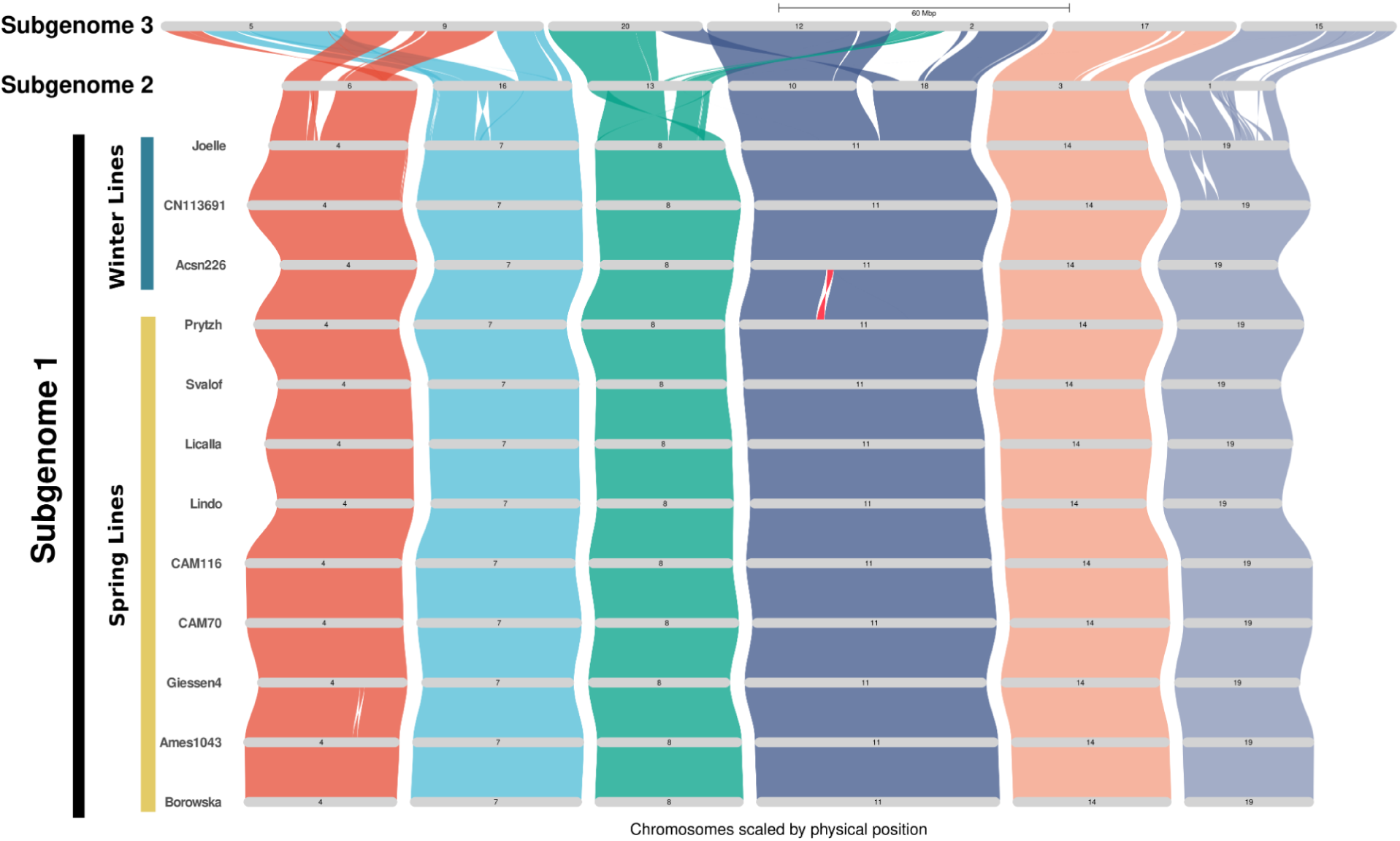
Riparian plot of genome synteny between subgenomes and between all lines for subgenome 1. Winter lines which require vernalization are marked by the blue bar, spring lines which do not require vernalization are marked by the yellow bar. The 100kb inversion of interest is highlighted in red.

### Acyl-lipid metabolism genes are highly conserved across lines but divergent between Camelina and Arabidopsis

*Camelina sativa* is an oilseed crop that has had a large expansion in the metabolic flux contributed to the production of fatty acids within the seed in comparison to the typical Brassicaceae. We thus proceeded to better understand if this is a property of the pangenome variation within the species or is possibly influenced by the deeper orthogroup variation across species. To do this, we investigated copy number change and the extent of pangenome variation in genes involved in acyl-lipid metabolism. Based on a curated list of lipid-associated *Arabidopsis thalian*a genes and the annotated enzyme codes and GO terms of Camelina genes, we identified a total of 1019 acyl-lipid related orthogroups, 215 of which did not have an Arabidopsis thaliana ortholog (Fig S6; Table S16).

We first assessed the extent of presence-absence and copy number variations among lines for lipid-associated orthogroups. We found that only 17% of acyl-lipid orthogroups exhibited presence-absence variation. Of the 804 orthogroups with Arabidopsis orthologs, 573 (71%) exhibit no variation in gene copy number across the twelve Camelina lines. Of those with no copy number variation, 507 (67%) orthogroups have the expected 3:1 copy number ratio between Camelina and Arabidopsis (Fig10; Table S17). This is a marked increase compared to the proportion of total orthogroups found in a 3:1 copy number ratio across all lines (43%; Fig10; Table S17). These results indicate a minimal contribution of copy number variation to acyl-lipid variation in Camelina.

Based on our observation of substantial orthogroup gain and loss between Arabidopsis and Camelina, we next queried whether these taxonomically restricted orthogroups contributed to acyl-lipid metabolism. We found that only four (0.4%) of lipid-associated orthogroups contained only Arabidopsis orthologs, far fewer than what was observed for all orthogroups (6%; Fig 10; Table S17), suggesting greater than average retention of lipid-associated genes. In contrast, the proportion of orthogroups that are Camelina specific was similar between lipid-related genes (21%) and all orthogroups (24%; Fig 10; Table S17). Thus, orthogroup variation across species contributes to acyl-lipid metabolism, either through the gain of orthogroups in Camelina or loss in Arabidopsis.

To determine if these taxonomically restricted orthogroups are potentially associated with specific shifts in lipid metabolism, we tested for over- and under-representation of specific Enzyme Codes among these Camelina-specific orthogroups using a Χ^2^ test. We identified 15 Enzyme Codes that significantly departed from the expected frequency. We identified codes representing 2-acylglycerol O-acyltransferase and diacylglycerol O-acyltransferase were over-represented (Table S18). Genes with these acyltransferase activities (DGAT1 and DGAT2) were previously associated with oil quality in *C. sativa*, suggesting these taxonomically restricted orthogroups are concentrated to specific aspects of lipid metabolism and may also play a role in pathways relevant to Camelina oil quality. Among the under-represented enzyme codes were monoglyceride lipase and Ketoacyl-CoA Synthase (Table S18).

**Fig 10.**
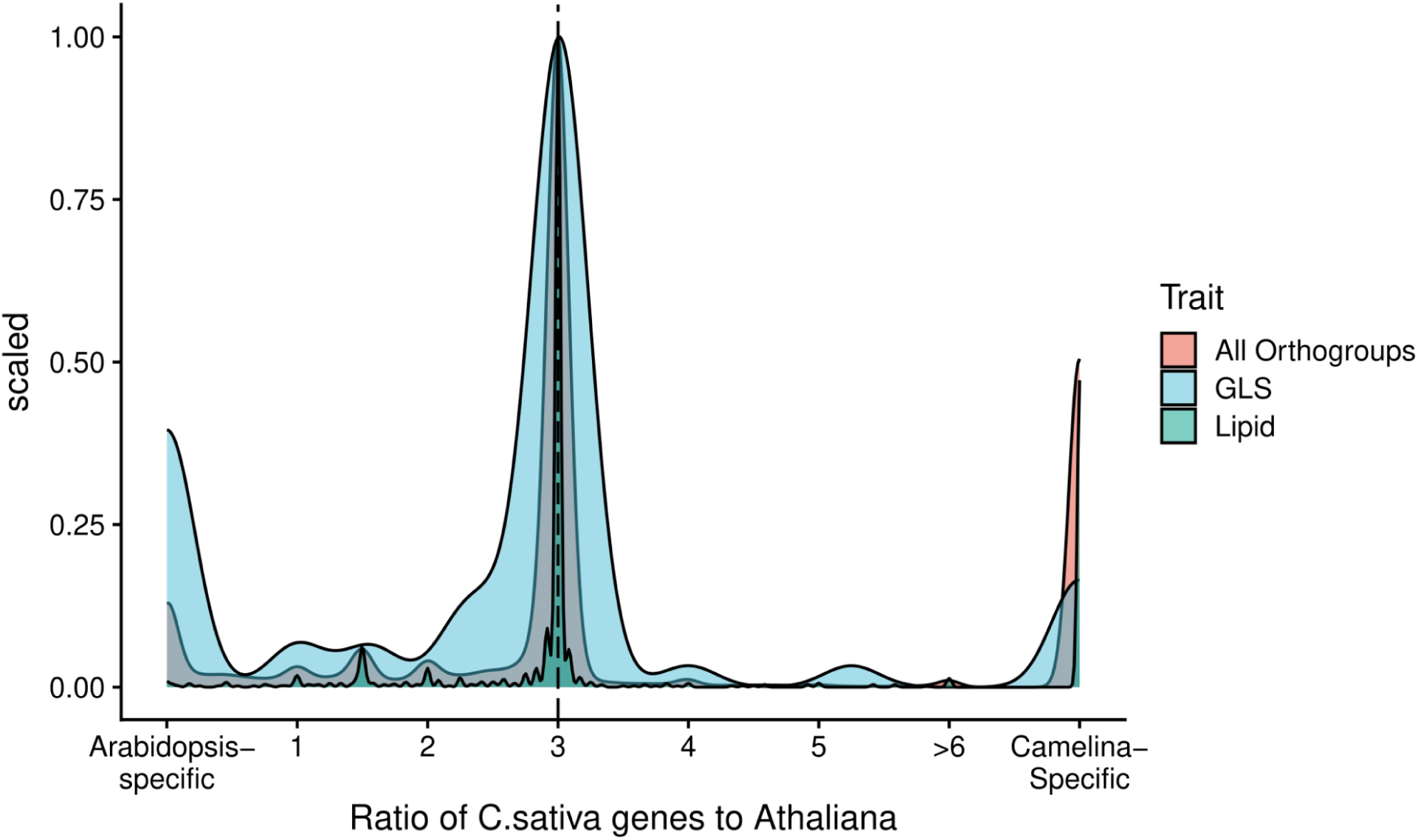
Density plot showing the ratio of *Camelina sativa* gene copy number to Arabidopsis copy number for all orthogroups (red), orthogroups of genes involved with indolic and aliphatic glucosinolate biosynthesis (blue), and orthogroups of genes involved in acyl-lipid metabolism (green).

Genes involved in the latter activity are involved in very-long-chain fatty acid formation and a gene from this enzyme class, Fatty Acid Elongase 1 is related to oil quality (Haslam and Kunst, 2013; Neumann et al. 2021).

While the bulk of acyl-lipid orthogroups appear to be highly stable in gene content across the lines, we noticed that the genes from novel orthogroups were enriched for pangenome variation. Genes in Camelina-specific orthogroups were more variable across the pangenome with a coefficient of variation of 1.28 versus only 0.086 for genes in orthogroups with an Arabidopsis ortholog. This suggests that these novel orthogroups may also contribute a significant portion of variation in lipid metabolism within the species.

### Novel gene gains and losses underlie glucosinolate pathway rewiring and change in chemotype between Camelina and Arabidopsis

Another key trait differentiating Camelina from Arabidopsis and other Brassicaceae is the specific glucosinolates accumulated. In other species like Brassica and Arabidopsis, glucosinolate and other specialized metabolites genes are noted for having pangenome variation at elevated rates with this variation causally controlling the glucosinolate biosynthesis (Kliebenstein et al. 2001; Kroyman, 2003; Chan et al. 2010; Hofberger et al. 2013; Barco and Clay, 2019; Katz et al. 2021). To test if this pattern of elevated pangenome variation is also true in Camelina, we phylogenetically analyzed homologs of known GLS identified through a BLAST search of our genomes using the pipeline blast-align-tree (https://github.com/steinbrennerlab/blast-align-tree/tree/main). The resulting gene trees were manually inspected to confidently identify orthologous and paralogous genes in Camelina based on phylogenetic placement relative to Arabidopsis. These genes were then assessed for copy number and presence-absence variation.

Globally we observed a greater proportion of genes exhibiting copy number variation in the glucosinolate pathway, demonstrated by the wider distribution of copy number ratio. The additional peaks at intermediate values in the copy number distribution show that there is also a greater fraction of genes with partial expansions or contractions in copy number relative to Arabidopsis (Fig 10; Table S17). As found for lipids, there is a large enrichment in Camelina-specific orthogroups for glucosinolates. In contrast to lipids, there is also a larger fraction of genes absent in Camelina as represented by the boost in Arabidopsis-specific orthogroups (Fig 10; Table S17). This suggests that glucosinolate variation is driven more by orthogroup variation across species and subgenomes rather than pangenomic variation among lines.

We queried the orthogroup variation to test if it is possible to explain the novel long-chain aliphatic glucosinolate phenotype in Camelina versus Arabidopsis. Interestingly, Camelina appears to have novel orthogroups in a number of steps associated with chain elongation including a gain of a novel MAM gene (Fig S7), two novel CYP79F genes (Fig S8), duplication of the long-chain FMO_GS-OX5_ and gain of a novel BCAT (Fig 11; Table S19). This suggests that the gain in long-chain glucosinolates required the gain of a number of new genes/steps.

Associated with the gain in long-chain glucosinolates is the loss in side-chain modification of aliphatic glucosinolates within Camlina. This is reflected by the loss of short-chain modification genes like CYP79F1, CYP79F2, FMOGS-OX1, FMOGS-OX3, FMOGS-OX4, AOP2, AOP3, and GS-OH (Fig 11;Table S19). Thus, the novel glucosinolate phenotype in Camelina involves major gains and losses in gene content across the entire pathway.

Camelina is also known to possess several indolic glucosinolates, which are derived from tryptophan rather than methionine. We investigated gene content variation of indolic glucosinolates biosynthesis genes. We observed that, like for aliphatic glucosinolates, the majority of variation is concentrated in the side-chain modification genes (Fig S9; Table S20). We confirmed previous studies which identified widespread loss of known CYP81F genes, except for CYP81F3, and a novel CYP81F gene previously called CYP81F6. Contrary to previous reports that *C. sativa* possesses 3 copies of CYP81F6, we found our lines have either one or two copies of CYP81F6. We additionally found the previously identified independent expansions of indole glucosinolate O-methyltransferase (IGMT) genes. Each Camelina line possesses four IGMT genes which are sister to IGMT1-IGMT4 from *Arabidopsis*.

In addition to biosynthesis, there are also significant changes to glucosinolate activation pathways (Fig 11; Table S19). In agreement with the absence of alkenyl glucosinolates in *C. sativa*, there is a complete loss of ESP. NSP5, which is likely the ancestral gene from which NSP1 to NSP4 derive, was found at the expected 1:3 ratio compared to Arabidopsis.

Meanwhile, all the rest of the NSPs appear to be lineage-specific expansions. Arabidopsis NSP1 to NSP4 are in a single clade that is sister to the clade containing all other Camelina NSP homologues. Within this, there was pangenomic variation with eight Camelina lines possessing six NSP homologs, two lines with seven homologs, and two with eight NSP homologs. The topology of the gene tree suggests independent expansions of an ancestral NSP gene in these two lineages.

**Fig 11.**
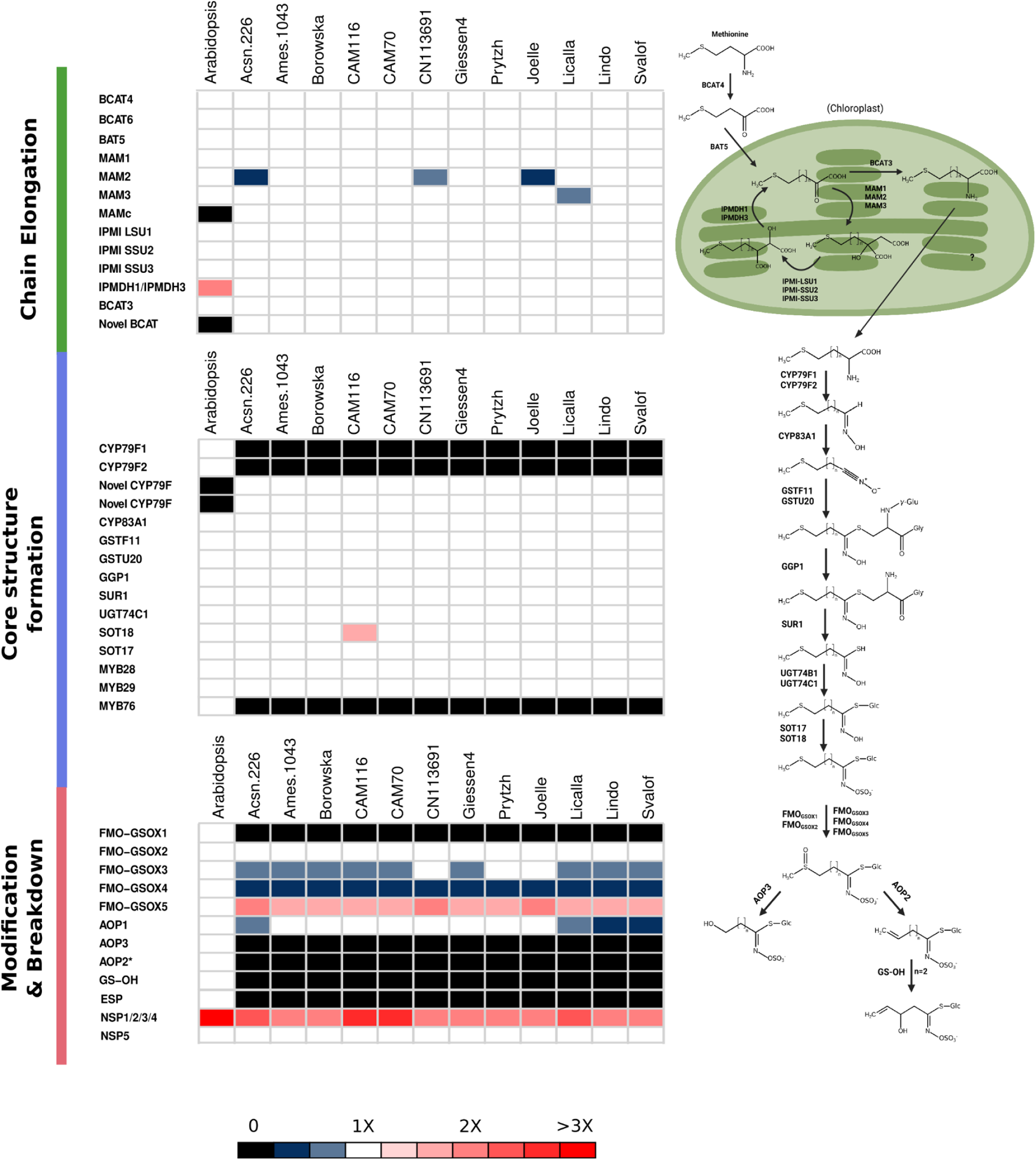
Varying copy number of homologs involved in the aliphatic glucosinolate pathway, relative to expected copy number in Arabidopsis (1) and our twelve Camelina genomes (3). We refer to the fourth MAM in Camelina as “MAMc” following Abrahams et al. (2020).

## Discussion

To explore the potential role of WGD in shaping genomic variation in recently formed polyploid species, we constructed twelve complete, chromosome-scale *Camelina sativa* genomes and conducted comparative pangenomic analyses. This effort represents one of the most complex genome structures to receive pangenomic resources and complements existing reference-quality assembly based pangenomes in polyploid species.

### Camelina sativa genome structure and gene content is highly conserved across accessions

Ancient whole-genome duplications have played a prominent role in eukaryotic evolution, but the consequences of WGD in recently formed polyploids are still unclear. Based on theory and recent data showing greater levels of pangenomic variation in polyploid species (Song et al. 2020; Hamala et al. 2023), we expected high levels of gene presence-absence variation (PAV) and structural variation (SV) in the Camelina pangenome. Our high-quality genomes assessed these expectations with the first detailed characterization of pangenomic variation in Camelina. In contrast to expectations, this revealed very low levels of gene PAV and SVs. The Camelina pangenome had relatively lower diversity, exhibiting fewer orthogroups affected by PAV, and less of the genomic sequence space affected by SVs in comparison to a polyploid from the same family, *Brassica napus* (Song et al. 2018). It also showed lower levels of PAV and SV than a more closely related diploid, *Arabidopsis thaliana,* further suggesting that WGD doesn’t increase the rate of pangenomic variation (Kang et al. 2023; Lian et al. 2024). The low rate of pangenome variation agrees with estimates of low of nucleotide and phenotypic diversity in *C. sativa* (Luo et al. 2019; Li et al. 2021; Brock et al. 2022b). In combination this suggests that WGDs do not automatically result in high levels of pangenomic variation.

The disparity in the pangenomes of Camelina and *B. napus* raises the question of what factors contribute to variation in pangenome size between polyploid lineages. Given that homoeologous exchanges were reported to contribute to PAV and SVs in natural and synthetic *B. napus* (Hurgobin et al. 2018; Bayer et al. 2022), factors affecting the degree of homoeologous recombination may explain the differences in pangenomic variation between the two species. In line with this, wheat, which is known to have almost complete suppression of homoeologous crossovers, exhibits similar levels of PAV to Camelina (Walkowiak et al. 2020). The paucity of homoeologous exchange events identified in our newly constructed *C. sativa* pangenome, on average only three HEs per lines affecting ∼0.03% of the genome, suggests it is more meiotically stable than *B. napus,* and may contribute to its smaller pangenome.

In all polyploids, establishing meiotic stability is critical for maintaining fitness and allopolyploids are known to achieve meiotic stability through the suppression of homoeologous recombination and associated exchanges (Gaeta and Pires, 2010; Lloyd and Bomblies, 2016; Cheron et al. 2023). This may suggest that selection for greater meiotic stability in allopolyploids can come at the cost of the pangenomic variation generated by homoeologous exchanges. Different allopolyploid species may adopt different balances between genomic variation and meiotic stability leading to across-species variation in pangenome sizes. For example, while natural *B. napus* is more meiotically stable than synthetic lines, it does not fully suppress homoeologous recombination like hexaploid wheat and therefore generates greater levels of homoeologous exchange (Grandont et al. 2014; Higgins et al. 2021; Sourdille and Jenczewski 2021). Future pangenomic studies in polyploid species can shed light on the factors influencing meiotic stability of allopolyploids and how this affects the relationship between WGD and pangenomic variation.

### Hybridization likely expanded gene content and functional potential in *Camelina sativa*

The presence of pangenome presence-absence variation within a species can expand to gene content variation across species due to drift during lineage sorting and adaptive gene loss leading to new species fixing for the absence of a gene. Several “super pangenome” studies measuring gene PAV between species support this expectation. Specifically querying species in the genera *Fragaria, Vitis, Eucalyptus,* and *Solanum* showed that they all exhibit high levels of species-level divergence in gene content (Qiao et al. 2021; Bozan et al. 2023; Li et al. 2023; Cochetel et al. 2023; Ferguson et al. 2024). This variation in gene or orthogroup variation between species means that allopolyploidy can increase gene content by merging different species genomes. Ultimately, this could expand the functional potential of an allopolyploid species. Our genomes support that this may have occurred in the Camelina allopolyploid event by showing that 19% of all Camelina genes were specific to a single subgenome. This is most likely representing gene content in the progenitor diploids as the recent age of the polyploid origin suggests that there hasn’t been sufficient time for the wide-spread, independent gene loss on multiple subgenomes necessary to create this post-polyploid formation. Similarly, all subgenomes showed a similar level of specific gene content regardless if they contributed to the first or second polyploid origin leading to the allopolyploid. The amount of subgenome-specific genes reported here in Camelina is consistent with reports in the *B. napus*, hexaploid wheat, and *Arabidopsis suecica* genomes (Chalhoub et al. 2014; IWGSC, 2018; Burns et al. 2021), suggesting the expansion of gene content from hybridization may be a common consequence of allopolyploidy.

This merger of genome functionality may specifically affect the resulting potential of an allopolyploid species. The subgenome-specific genes showed enrichment for specific functions that differed based on the ancestral diploid being queried. These included enrichments in functional terms, such as auxin response, signal transduction, defense response, and tRNA aminoacylation, which has been shown to be involved in plant defense (Soprano et al. 2018) and may play even broader functional roles through potential non-canonical function (Saga et al. 2020). This suggests the potential for gene content divergence between diploids, driven by neutral drift or differential adaptation, to affect biotic and abiotic stress responses that upon whole genome merger in an allopolyploid could increase the functionality of the resulting polyploid. Future work will have to assess the breadth of how combining subgenome-specific genes may create a benefit during polyploid formation and how this compares to the evolutionary potential created by gene duplication.

### Lineage specific genes may contribute to Camelina’s metabolic novelty

Given the potential contribution of subgenomic specific genes in contributing to novelty in this system, we queried how deeper lineage specificity may contribute. Previous work had shown that pangenomic and super-pangenomic variation has been associated with metabolic variation and biotic and abiotic adaptations (Hurgobin et al. 2018; Katz et al. 2021). Testing how orthogroup changes across phylogenetic levels linked to variation in key metabolic pathways identified potential connections to metabolic novelty. Our investigation of the acyl-lipid and glucosinolate pathways found novel orthogroups present only in Camelina and not Arabidopsis could explain unique metabolic profiles in Camelina. For lipid metabolism, Camelina appears to have expanded its gene content, with 21% of orthogroups being Camelina specific and only 0.4% Arabidopsis-specific. We specifically found enrichment of Camelina-specific genes for diacylglycerol O-acyltransferase activity. Genes of this enzyme class, such as DGAT1, DGAT2, and DGAT3, have been previously implicated in oil quality traits, (Lager et al. 2020; Gao et al. 2021; Lee et al. 2022) suggests this expansion contributes to key oil quality traits that make Camelina a favorable biofuel crop.

In the glucosinolate pathway, gain of novel genes in Camelina appear to have contributed to the evolution of their unique long-chain aliphatic glucosinolates. These gains were paired with the loss of side-chain modification genes known to play a key role in Arabidopsis adaptive evolution (Zust et al. 2012; Brachi et al. 2015; Kerwin et al. 2015; Kerwin et al. 2017). These modification enzymes are known to act on short-chain aliphatic glucosinolate. This may suggest that long-chain glucosinolate capacity evolved first and the metabolic shift nullified the function of side-chain modification genes, resulting in gene loss. Critically, many of these shifts in gene content like gain of MAMc and novel CYP79F homologs were found to be shared with *Capsella rubella*, a closely related species with a similar chemotype (Czerniawski et al. 2021). This suggests the metabolic shift in Camelina occurred further back in the evolutionary lineage and may explain the general lack of subgenome-level variation in the glucosinolate pathway as these genomic and metabolic changes existed in the common ancestor of the diploid progenitors. These results highlight that candidates for crop improvement are likely to exist beyond Arabidopsis orthologs. Future mechanistic work is required to test these hypothesis and deeper phylogenetic analysis is required to assess when these novel processes evolved in the lineage leading to Camelina or if they were lost in the Arabidopsis lineages.

## Acknowledgements

The work (proposal: Award DOI 10.46936/10.25585/60001293) conducted by the U.S. Department of Energy Joint Genome Institute (https://ror.org/04xm1d337), a DOE Office of Science User Facility, is supported by the Office of Science of the U.S. Department of Energy operated under Contract No. DE-AC02-05CH11231. This work was also supported by NSF-IOS PRFB 2208944 to K.A.B, USDA 2019-05709 and NSF MCB 1906486 to D.J.K, NSF-IOS PRFB 2109178 to J.R.B, and NSF-PGRP 2029959 and DOE-BER DE-SC0022987 to P.P.E.

## Data availability

Reference genome assembly and annotation files of Acsn-226 (v 1.1), Ames 1043 (v 1.1), Borowska (v 1.1), CAM116 (v 1.1.), CAM 70 (v 1.1.), CN113691 (v 1.1), Giessen4 (v 1.1), Joelle (v 1.1), Licalla (v 1.1), Lindo (v 1.1), Prytzh (v 1.1), and Svalof (v 1.1) genomes are available at https://phytozome-next.jgi.doe.gov/. All raw sequence reads have been deposited in the NCBI SRA database under BioProject accessions PRJNA1143072, PRJNA1143073, PRJNA1143074,PRJNA1143075, PRJNA1143076, PRJNA1143077, PRJNA1143078, PRJNA1143079, PRJNA1143080, and PRJNA1143081. Supplementary tables S1-S23 are uploaded to the GSA Figshare portal. Other intermediate files are provided at the Zenodo archive https://doi.org/10.5281/zenodo.13158027. Code to recreate main results and figures are available at https://github.com/KevinABird/CsativaPangenome/

## Methods

### Plant Material, DNA and RNA extraction

For genomic sequencing, plants were grown in a Michigan State University greenhouse in East Lansing, MI. Eight week old plants were dark treated for 48 hours prior to sampling and the youngest leaves were sampled, flash frozen in liquid nitrogen, and sent to the Arizona Genome Institute (AGI) for high-molecular weight genomic DNA extraction.

For RNAseq and Iso-Seq, plants were grown under 23°C : 20°C, 12h : 12h day : night cycles in a growth chamber. Several Leaf tissue across the diurnal cycle was sampled at ZT0, ZT6, ZT12, and ZT18,where ZT refers to Zeitgeber Time and indicates the number of hours since the last dark-to-light transition. Stem, flower, immature fruit, cauline leaves, and leaves 48 hours post treatment with 5mM silver nitrate, were collected within one hour of 11am (ZT3) and flash frozen in liquid nitrogen. RNA was extracted from frozen tissues using the Qiagen RNeasy Plant kit (Qiagen, Hilden, Germany). RNA purity was assessed using a Qubit (ThermoFisher, Waltham, MA). Samples from leaf tissue (diurnal samples, cauline leaves, silver-nitrate treatment), and other tissues (stem, flower, fruit) were combined to make two separate pooled RNA samples per accession.

### Genomic sequence and genome assembly

#### Sequencing and library construction

The Camelina PacBio HiFi libraries were constructed using Circular Consensus Sequencing (CCS) mode. DNA shearing was accomplished using a Diagenode Megaruptor 3 instrument. We constructed libraries using SMRTbell Template Prep Kit 2.0, which were then tightly sized on a SAGE ELF instrument (1-18kb) to a final library average insert size of 20kb. The Dovetail OmniC library for Prytzh was built using standard protocols (Dovetail Omni-C kit Catalog #21005). Illumina libraries were built using standard protocols (Illumina TruSeq PCRfree Catalog #20015962).

We sequenced the Camelina genomes using a whole genome shotgun sequencing strategy and standard sequencing protocols. Sequencing reads were collected using PACBIO and Illumina platforms. PACBIO and Illumina reads were sequenced at the HudsonAlpha Institute in Huntsville, Alabama. PACBIO reads were sequenced using the SEQUEL II platform and Illumina reads were sequenced using the Illumina NovoSeq 6000 platform.

For the PACBIO sequencing, each genotype was sequenced using 1-3 SMRT cells using V2 chemistry, producing total raw sequence yields of 15.6-66.8 Gb and total coverage of 19.54x-83.51x (Table S17). We also sequenced a 400bp insert 2×150bp Illumina fragment library for each genotype (48.4x-65.1x), and for Prytzh, we sequenced one 2×150 HiC Library (Table S21). Prior to assembly, we screened Illumina fragment reads for phix contamination. Reads composed of >95% simple sequence were removed. Illumina reads <50bp after trimming for adapter and quality (q<20) were removed.

#### Genome assemblies and construction of pseudomolecule chromosomes

The Camelina assemblies were generated by assembling PACBIO CCS reads (1,002,068-3,177,007 reads, 19.54x-83.51x coverage, 14,485-25,809 bp average read size; Table S18) using HifFiAsm (v0.16.1-r375, Cheng et al. 2022; HiC integration [-h1 and -h2] used for Pryzth v1.0). The output from the assemblers were subsequently polished using RACON (v1.4.10, Vaser et al. 2017), producing initial assemblies consisting of 793-1793 scaffolds (contigs), with contig N50s of 11.6-30.4 Mb and total genome sizes of 669.1-744.2 Mb (Table S22).

For Prytzh, Hi-C Illumina reads were separately aligned to the contigs with Juicer (v1.5; Durand et al. 2016), and chromosome scale scaffolding was performed with 3D-DNA (v.180922; Dudchenko et al. 2017). The HiC data was used to search for contig breaks, where there is an abrupt change in linkage group within a contig, and no misjoins were identified. The contigs were then oriented, ordered, and joined together into 20 chromosomes using the HiC data. A total of 52 joins were applied to the Prytzh assembly.

For the other 11 assemblies, 20,563 syntenic markers generated from the Pryzth v1.0 assembly (see below) were used to identify contig breaks in the polished assemblies. Five assemblies had zero misjoins, and six assemblies required 1-3 contig breaks (Table S22). The sytentic markers from Pryzth v1.0 were then used to orient, order, and join contigs into 20 chromosomes. A total of 7-78 joins (Table S22) were made during this process. Nine assemblies included 1-3 (Table S22) additional scaffolds not incorporated into one of the chromosome-scale scaffolds.

For all 12 assemblies, after forming the chromosomes, it was observed that some small (<20Kb) redundant sequences were present on adjacent contig ends within chromosomes. To resolve this issue, adjacent contig ends were aligned to one another using BLAT (Kent, 2002), and duplicate sequences were collapsed to close the gap between them. A summary of these collapsed adjacent contigs can be found in Table S22. Each chromosome join is padded with 10,000 Ns. Contigs terminating in significant telomeric sequence were identified using the (TTTAGGG)_n_ repeat, and care was taken to make sure that they were properly oriented in the production assembly. Scaffolds that were not anchored in a chromosome were classified into bins depending on sequence content. Contamination was identified using blastn against the NCBI non-redundant nucleotide collection (NR/NT) and blastx using a set of known microbial proteins. Additional scaffolds were classified in the released assemblies as repetitive (>95% masked with 24mers that occur more than four times in the genome), redundant (unanchored sequence with >95% identity and >95% coverage within a chromosome), chloroplast, and mitochondria.

In three assemblies, 1-3 large homopolymer regions were identified within contigs (Table S18). The CCS reads were aligned using minimap2 (v2.24, Li 2021) to the chromosomes of these assemblies and revealed that these regions had very low depth and were likely sequencing artifacts. These regions were removed and replaced by gaps of 10,000Ns.

Finally, homozygous SNPs and INDELs were corrected using ∼48x-65x of Illumina reads (2×150, 400bp insert) by aligning the reads using bwa mem (v2.2.1, Li and Durbin 2009) and identifying homozygous SNPs and INDELs with the GATK’s UnifiedGenotyper tool (v3.7, McKenna et al. 2010). A total of 3-276 homozygous SNPs and 327-2542 homozygous INDELs were corrected in our releases (Table S22). The final versions of the assemblies contain 597.0-656.3 Mb of sequence, consisting of 26-95 contigs with contig N50s of 13.3-31.3 Mb and a total of 99.82%-100% of assembled bases in chromosomes (Table S22).

From the polished Pryzth assembly, 20,563 unique, non-repetitive, non-overlapping 1 kb syntentic markers were generated to be used during the assembly of the remaining 11 C.sativa genome assemblies. For each assembly, the syntenic markers were aligned using BLAT (Kent 2002) to contigs in order to order and orient contigs in pseudomolecules. This order and orientation was confirmed and breaks were made as necessary.

Completeness of the euchromatic portion of the assemblies was assessed using 82,471 primary transcripts from the C.sativa var. DH55 genome assembly (Kagale et al. 2014). The aim of this analysis is to obtain a measure of completeness of the assembly, rather than a comprehensive examination of gene space. The transcripts were aligned to each assembly using BLAT (Kent 2002) and alignments >=90% base pair identity and >=85% coverage were retained. The screened alignments indicate that 99.21%-99.33% of the transcripts aligned to our assemblies (Table S22).

### RNA sequencing and Genome Annotation

#### RNA Sequencing

For PacBio-based Iso-Seq RNA sequencing, 500ng of total RNA is used as input for each library (Table S23). Full-length cDNA is synthesized using template switching technology with NEBNext Single Cell/Low Input cDNA Synthesis & Amplification Module kit (Catalog #E6421). The first-strand cDNA is amplified and multiplexed with NEBNext High-Fidelity 2x PCR Master Mix (Catalog #M0541) using barcoded cDNA PCR primers. The amplified cDNA (11-14 cycles) is purified using 1x AMPure PB beads (PacBio Catalog #100-265-900) for non-size selection or BluePippin (Sage Science, Beverly, MA) for 2-10 kb size selection and liked sizes are pooled at the equimolar ratios in a designated Degree-of-Pool in the worksheet using PacBio Multiplexing Calculator (PacBio, Benlo Park, CA). The pooled samples are end-repaired, A-tailed and ligated with overhang non-barcoded adaptors using SMRTbell Express 2.0 kit (PacBio Catalog #100-938-900). PacBio Sequencing primer is then annealed to the SMRTbell template library and sequencing polymerase was bound to them using Sequel II Binding kit 2.0 (PacBio Catalog #101-789-500). The prepared SMRTbell template libraries are then sequenced on a PacBio Sequel IIe sequencer using SMRT Link 10.2, sample dependent sequencing primer, 8M v1 SMRT cells, and Version 2.0 sequencing chemistry with 1×1800 sequencing movie run times (PacBio, Menlo Park, CA).

For Illumina-based RNAseq RNA sequencing, plate-based RNA sample prep is performed on the PerkinElmer Sciclone NGS robotic liquid handling system (Revvity, Waltham, MA) using Illumina’s TruSeq Stranded mRNA HT sample prep kit (Catalog #20020595) utilizing poly-A selection of mRNA following the protocol outlined by Illumina in their user guide: https://support.illumina.com/sequencing/sequencing_kits/truseq-stranded-mrna.html, and with the following conditions: total RNA starting material is 1000 ng per sample and 8 cycles of PCR is used for library amplification. The prepared libraries (Table S23) are quantified using KAPA Biosystems’ next-generation sequencing library qPCR kit (Roche Catalog #07980140001) and run on a Roche LightCycler 480 real-time PCR instrument (Rocche Diagnostics, Indianapolis, IN). Sequencing of the flowcell is performed on the Illumina NovaSeq sequencer using NovaSeq XP V1.5 reagent kits (Illumina, San Diego, CA), S4 flowcell, following a 2×151 indexed run recipe.

#### Genome Annotation

Genome annotation was performed in two rounds. In the first round, each of the twelve assemblies was annotated individually. Subsequently, the annotations from each assembly were used to expand and improve the annotations of all the assemblies in the pangenome.

During the first round, for each genotype, we generated transcript assemblies from 2×150bp Illumina RNA-seq reads (Table S19) using PERTRAN (described in detail in Lovel et al. 2018). PERTRAN uses GSNAP (Wu et al. 2010) for genome-guided transcriptome assembly, then validates, realigns, and corrects alignments, and builds a splice alignment graph. We used PacBio Iso-Seq CCS reads to obtain putative full-length transcripts by aligning the CCS reads to the genome with GMAP (Wu et al. 2010), correcting small indels in splice junctions, and clustering identical multi-exon alignments and single-exon alignments that had at least 95% overlap. PASA (Haas et al. 2003) was then used to construct processed transcript assemblies using the RNA-seq transcript assemblies, corrected CCS clusters, and Sanger ESTs.

To repeat-soft-mask the assemblies, we first generated a *Camelina* species-level repeat library from *de novo* repeats predicted by RepeatModeler (Smit, 2008-2015) using the *Camelina sativa* CAM116 v1.0 genome assembly. We used InterProScan (Jones et al. 2014), including the Pfam (Mistry et al. 2021) and PANTHER (Mi et al. 2019) databases, for functional analysis of the predicted repeats. Any repeats with significant similarity to protein-coding domains were removed from the repeat library. We then used the species-specific repeat library to soft-mask each assembly using RepeatMasker (Smit et al. 2013-2015).

Putative gene loci were identified using alignments of processed transcript assemblies and/or alignments from EXONERATE (Slater, 2005) of proteins from *Arabidopsis thaliana*, soybean, poplar, cotton, rice, sorghum, peach, citrus, *Aquilegia*, tomato, grape, *Eutrema salsugineum*, *Nymphaea colorata*, *Amborella trichopoda*, *Cucurbita maxim*, *Capsella rubella*, and Swiss_Prot release 2021_03 of eukaryote proteomes to repeat-repeat-masked assemblies. The putative loci were extended up to 2kb in both directions of alignments unless they extended into another locus on the same strand. To predict gene models at each locus, we used homology-based predictors, FGENESH+ (Salamov et al. 2000), FGENESH_EST (similar to FGENESH+, but it uses ESTs to compute splice site and intron input rather than protein/translated ORF), EXONERATE, PASA assembly ORFs (in-house homology constrained ORF finder), and AUGUSTUS (Stanke et al 2006) that was trained on the high confidence PASA assembly ORFs and with intron hints from short read alignments. The best-scored prediction for each locus was selected using multiple factors, including EST and protein support and overlap with repeats. We improved the selected gene predictions using PASA by adding UTRs, splicing correction, and incorporating alternative transcripts.

The above-mentioned proteomes were used to obtain Cscores and protein coverage for the PASA-improved gene model proteins. We used Cscore, protein coverage, EST coverage, and CDS overlap with repeats to select transcripts. Transcripts were selected when their Cscore and protein coverage >= 0.5 or if covered by ESTs. If a gene model had CDS overlapped by repeats by more than 20%, then we required a Cscore >= 0.9 and >= 70% homology coverage. We performed Pfam (Mistry et al. 2021) analysis on the gene models and filtered out any where proteins had >30% overlap with Pfam TE domains.

During the second round of annotation, we hard-masked each genome using their own high-confidence gene models. We used BLASTX and EXONERATE to compare each hard-masked genome to all the high-confidence peptide predictions from the other assemblies and used the results to make EXONERATE gene predictions. BLASTP was used to score these new gene predictions using homology proteomes. New models (a) replaced models from the first-round if they had higher homology support and were not contradicted by transcriptome evidence or (b) were added if there were no first-round gene models at the locus.

We filtered out gene models if they were incomplete gene models, had low homology support without full transcript-support, were short single exons (<450bp CDS) without protein domains nor good expression, or were repetitive gene models without strong homology support. [I will add the citations to the References section once I verify formatting]

Transposable elements (TEs) were annotated in the twelve *C. sativa* genomes in order to understand patterns of TE accumulation across these lines. Annotation was done with the Extensive De-Novo TE Annotator (EDTA) v2.2.0 (Ou et al. 2019; Ou et al. 2022) for each individual genome following methods outlined in Kang et al. 2023 using the genome assembly and coding sequences for each accession. EDTA parameters ‘--species others, --sensitive 1, --anno 1, --step all’ were used to generate de novo TE libraries. In addition, the flag ‘--curatedlib’ was called, providing the manually curated *A. thaliana* TE library available through the EDTA GitHub at https://github.com/oushujun/EDTA/tree/master/database. After manual inspection of each annotation, we proceeded with the panEDTA workflow to reannotate each genome provided the annotations of all others, thus creating the pan-TE library.

### GENESPACE Comparative Genomics

Syntenic orthologues and paralogues Arabidopsis thaliana and the three subgenomes of all twelve Camelina sativa genomes were inferred via GENESPACE (v.1.2.3, Lovell et al. 2022) pipeline using default parameters. GENESPACE compares protein similarity scores into syntenic blocks using MCScanX and uses Orthofinder to search for orthologues/paralogues within synteny constrained blocks. Ortholog information is projected between reference genomes.

### Gene Presence-Absence Variation

Gene Presence-Absence variation was assessed based on the hierarchical orthogroup membership of genes as reported in the combBed.txt output file from GENESPACE. Subgenomes were merged and an orthogroup was classified as present if an accession had at least one gene in that orthogroup and absent otherwise. We defined core orthogroups as those present in all twelve genomes, dispensable if present in between two and eleven lines, and unique if present in only one accession. For pangenome modelling, we sampled the core and pangenome size for all combinations of *n* genomes for *n*=2 to *n*=12. These values were plotted and a curve was fit with the geom_smooth() function using the loess method. To characterize deeper phylogenetic variation in orthogroup content, we identified orthogroups present or absent in Arabidopsis and the three Camelina subgenomes. The combination of orthogroup presence/absence across species and subgenomes was collated and visualized using the SankyeNetwork function in the R pacakge NetworkD3. Gene Ontology enrichment of subgenome-specific orthogroups was carried out using TopGO v2.56 (Alexa and Rahnenfuhrer, 2024).

### Structural Variants and Pangenome graph construction

To identify the large structural rearrangements (inversions, translocations, deletions, and duplications) and local variations (insertions and deletions), each genome was aligned to the Prytzh assembly using minimap2 (v2.26; Li 2021) with asm5 preset options. The resulting alignments were filtered in R to retain segments ≥ 10 kb in length and with ≥ 95% sequence identity and SyRI (v1.6.3; Goel et al. 2019) was used to classify these alignments as syntenic regions or structural rearrangements. Whole genome alignments were also used to construct a subgenome-aware pangenome graph with Minigraph-Cactus and default settings (v2.8.2; Hickey et al. 2023). Chromosome-level graphs were built using the Prytzh assembly as the primary reference and subsequently merged using vg (v1.54.0; Garrison et al. 2018). The degree of sequence sharing across genomes was assessed with Panacus (v0.2.3; Parmigiani et al. 2024)

### Acyl-lipid metabolism gene analysis

We identified a curated set of Arabidopsis genes involved with Acyl lipid metabolism (Li-Beisson et al. 2013). To identify Camelina genes without Arabidopsis orthologs but involved with Acyl-lipid metabolism we queried the functional annotations of our Camelina genomes for genes assigned an Enzyme Code or GO term associated with a gene from the curated Arabidopsis acyl-lipid metabolism list. These lists were manually curated, combined, and queried against members of the identified hierarchical orthogroups to create a list of unique hierarchical orthogroups representing the genes involved in acyl-lipid metabolism.

### Glucosinolate pathway gene analysis

A list of high-confidence genes involved in aliphatic and indolic glucosinolate biosynthesis was constructed based on Sonderby et al. (2010) and supplemented by reports of glucosinolate activation pathways for aliphatic glucosinolates (Burow et al. 2007) and side-chain modification genes for indolic glucosinolates (Czerniawski et al. 2021). We identified homologs of these known Arabidopsis glucosniolate genes using the BLAST-align-tree pipeline (https://github.com/steinbrennerlab/blast-align-tree/tree/main). This pipeline takes a user-selected query gene sequence and searches for a specified number of top BLAST hits in user-curated BLAST databases. These sequences are merged, aligned with ClustalOmege, and a phylogeny is made from the MSA using FastTree. Resulting phylogenies were manually inspected to identify the most appropriate outgroup, if the phylogenetic topology showed that Camelina genes were descendent to the specified Arabidopsis gene, the Camelina genes were considered orthologous. If Camelina genes formed a clade that was sister to a specified Arabidopsis gene, they were classified as novel paralogs. Phylogenetic trees for CYP79F and MAM genes were annotated and visualized using the interactive Tree of Life (iTOL) online tool (Letunic et al. 2021).

